# Computational Intractability Law Molds the Topology of Biological Networks

**DOI:** 10.1101/510495

**Authors:** Ali A Atiia, Corbin Hopper, Katsumi Inoue, Silvia Vidal, Jérôme Waldispühl

## Abstract

Virtually all molecular interaction networks (MINs), irrespective of organism or physiological context, have a majority of loosely-connected ‘leaf’ genes interacting with at most 1-3 genes, and a minority of highly-connected ‘hub’ genes interacting with at least 10 or more other genes. Previous reports proposed adaptive and non-adaptive hypotheses describing sufficient but not necessary conditions for the origin of this majority-leaves minority-hubs (mLmH) topology. We modeled the evolution of MINs as a computational optimization problem which describes the cost of conserving, deleting or mutating existing genes so as to maximize (minimize) the overall number of beneficial (damaging) interactions network-wide. The model 1) provides sufficient and, assuming 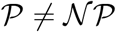, necessary conditions for the emergence of mLmH as an software adaptation to circumvent computational intractability, 2) predicts the percentage number of genes having *d* interacting partners, and 3) when employed as a fitness function in an evolutionary algorithm, produces mLmH-possessing synthetic networks whose degree distributions match those of equal-size MINs.

**Author Summary:** Our results indicate that the topology of molecular interaction networks is a selected-for software adaptation that minimizes the evolutionary cost of re-wiring the network in response to an evolutionary pressure to conserve, delete or mutate existing genes and interactions.

## 1 Introduction

Molecular interaction networks (MINs) are typically represented as graphs where nodes represent proteins, nucleic acids, or metabolites and edges represent physical or functional. By ‘functional’ we refer to the indirect effect that some gene *g_i_* has on another *g_j_*. None of the edges in the networks featured in this study represents a functional relationship, rather each edge represents a direct and physical interaction between *g_i_* and *g_j_*. interactions. The steady increase in scale [1] and resolution [2] of experimentally validated interactions has not been matched with theoretical progress towards deciphering the underlying evolutionary forces shaping the structure of MINs. Earlier studies aimed to empirically show that MINs possess a certain property (say, small-world connectivity or power-law-fitted degree distribution), and subsequently advocate for the existence of a universal design principle behind it. However, the robustness of the observations and conclusions of these studies [3, 4] has been questioned [5, 6, 7, 8], and so too [9, 10] has the validity of the design principles [11, 12] they inspired. Assuming those properties were indeed real, the universality of proposed design principles around them would still need to be justified. The statistical support of a property “is no evidence of universality without a concrete underlying theory to support it” [13]. Furthermore, a universal theory “must facilitate the inclusion of domain mechanisms and details” yet there is a sharp disconnect between current hypotheses’ high abstractions and actual functional aspects of evolving biolo*g_i_*cal systems [14]. A ‘software’ trait relates to *relationships* between genes (connectivity, community clustering, co-expression etc) rather than to their molecular (’hardware’) properties (e.g. their amino acid sequence or 3D conformation). The assumption that natural selection can lead to the emergence of advantageous network-level software traits has itself been challenged by the nonadaptive hypothesis: the topology of MINs could be an indirect result of selection pressure on other traits [15] or a mere byproduct of non-adaptive evolutionary forces such as mutation and genetic drift [16, 17]. Both the adaptive and non-adaptive hypotheses present sufficient but not necessary conditions for the emergence of network traits and therefore one cannot objectively rule out the plausibility of the other.

## 2 Perspective

A fundamental property in MINs is the majority-leaves minority-hubs (mLmH) topology^1^ whereby an overwhelming ~80% majority of leaf genes interact with at most 1-3 other genes, and an elite ~6% minority of hub genes interact with at least 10 other genes^2^. mLmH is observed in virtually all MINs irrespective of organism or physiological context as shown in Figure 1. Here we investigate the mLmH topology from a computational complexity perspective and derive necessary and sufficient conditions for its emergence. We assume that the organism is under some evolutionary pressure to change. Under this hypothetical scenario, we assume it critical for the system, in some regulatory state at some particular point in evolutionary time, that some genes be promoted and some be inhibited. An interaction is deemed beneficial if its promotional or inhibitory towards a gene that should indeed be promoted or inhibited respectively. Similarly, an interaction is deemed damaging if it is promotional or inhibitory towards a gene that should rather be inhibited or promoted, respectively. We assign a benefit (damage) score for each gene *g_i_* as the sum of beneficial (damaging) interactions that *g_i_* is *projecting onto* or *attracting from* other genes in the network. If gene *g_i_* promotes or inhibits *g_j_*, we say that *g_i_* is projecting an interaction onto *g_j_*, and *g_j_* is attracting an interaction from *g_i_*. The benefit or damage scores of both *g_i_* and *g_j_* is incremented by some some *ρ* ∈ ℝ If such an interaction is beneficial or damaging, respectively. *ρ* signifies the potency of such an interaction. A beneficial/damaging interaction therefore adds *ρ* to the total benefit or damage score of both the source and target genes. A gene may for example be projecting very few beneficial interactions while attracting many damaging interactions (making it more of a liability) and vice versa. Hence the benefit/damage scoring is not affected by how skewed the in-versus out-degree distribution of a gene is. Because we have no knowledge of what *ρ* is (i.e. measure of interaction potency) at a global scale in MINs, we use *ρ* = 1 for all interactions in this study.

**Figure 1:**
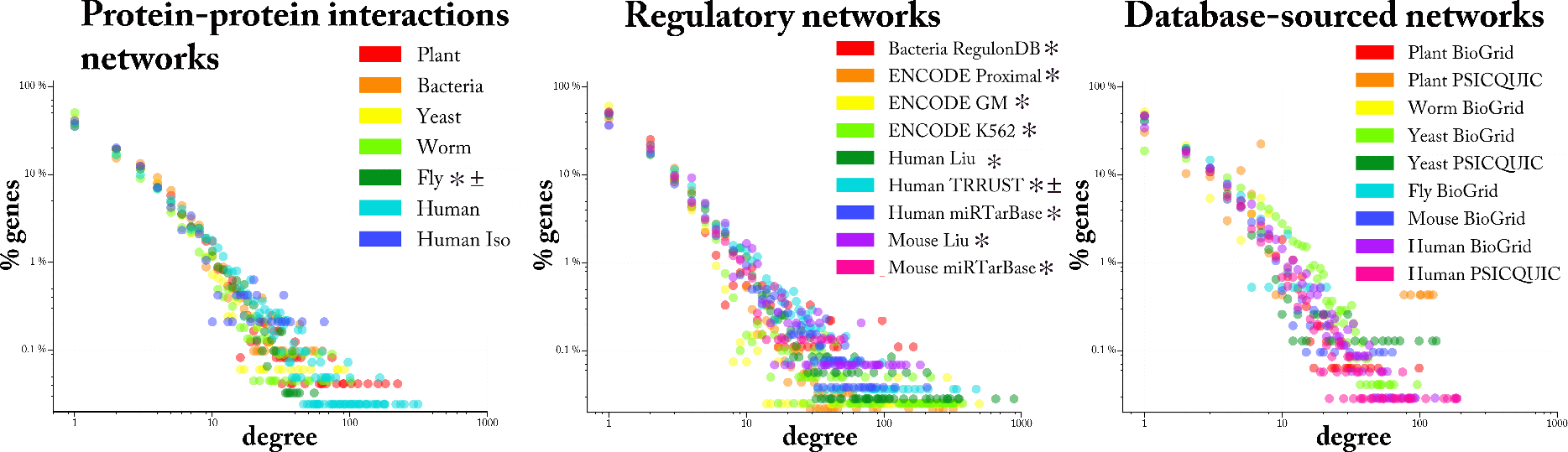
The majority-leaves minority-hubs (mLmH) topology in biological networks. Each dot represents the percentage of genes in the network having a given degree (number of interacting partners). On average, an overwhelming majority (~80%) of leaf genes interact with at most 1-3 other genes, and a small elite minority (~6%) of hub genes interact with at least 10 other genes. All networks originate from experimental procedures (i.e. none contains *in-* silico-inferred interactions), and all interactions are direct and physical. PPI and Regulatory networks are obtained from large-scale experimental studies reported in a single source in the literature, while interactions in database-sourced networks may have originated from more than one source (see SI 2 for details and literature references of each networks). Directed and signed networks are marked with * and ± respectively. The direction and sign were assigned at random (coin flip) in undirected/unsigned networks; some interactions in TRRUST are unsigned and hence were also assigned as random. Data and source code pertaining of all networks are publicly available in [39].

Given the benefit and damage scores of genes under the current evolutionary pressure scenario, how hard of a computational problem would it be to determine the optimal immediate “next-move” for the system, i.e. which genes to conserve, mutate or delete, such that the overall total number of beneficial interactions is maximal *while* the total number of damaging interactions is minimal to a threshold? We refer to this optimization question as the network evolution problem (NEP).

## 3 The Network Evolution Problem

The nodes in a MIN correspond to a set of genes *G* = {*g*_1_, *g*_2_, …, *g_n_*}, and the directed and signed edges represent interactions. The direction of an edge between *g_j_* and *g_k_* denotes which of the two genes is the source and which is the target. The nodes in the hypothetical MIN shown Figure 2 (A), left, represent 7 genes, with promotional and inhibitory interactions denoted by arrow- and bar-terminated edges, respectively. A hypothetical ‘Oracle advice’ (OA) on all or some of the genes simulates the evolutionary pressure on the network. The OA is represented as a ternary sequence *A* = (*a*_1_,*a*_2_, …, *a_n_*) where *a_j_* = +1 or *a_j_* = −1 indicates that the Oracle says *g_j_* should be promoted or inhibited, respectively. *a_j_* = 0 implies the Oracle is neutral towards *g_j_*. In Figure 2 (A) example, the OA is *A* = (0, +1, −1, +1, 0, 0, −1), meaning the Oracle says genes *g*_2_,*g*_4_ should be promoted (notice *a*_2_ = *a*_4_ = +1) while *g*_3_,*g*_7_ should be inhibited (*a*_3_ = *a*_7_ = −1). The Oracle has no opinion towards genes *g*_1_,*g*_5_ and *g*_6_ (*a*_1_ = *a*_5_ = *g*_6_ = 0) in this example.

**Figure 2:**
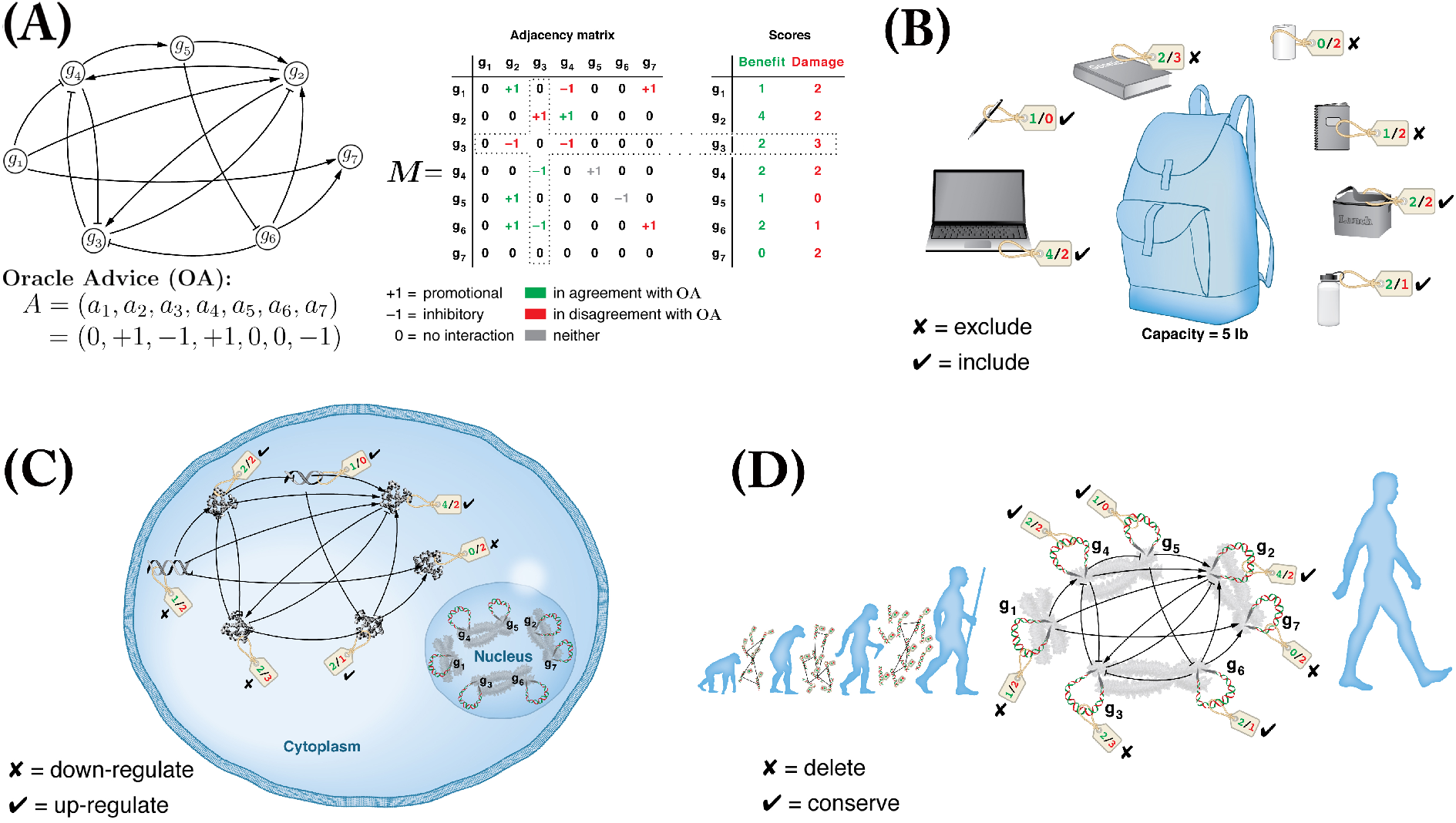
The network evolution problem (NEP). **(A)** Left: an example network of seven genes *G* = *g*_1_,*g*_2_,…,*g*_7_ that is under some hypothetical evolutionary pressure described by an Oracle advice (OA) *A* = (0, +1, −1, +1, 0, 0, −1) indicating that genes *g*_2_, *g*_4_ and *g*_3_, *g*_7_ should be promoted and inhibited respectively (notice +1 and −1 values in OA; *a_i_* ∈ *A* is the Oracle’s opinion on gene *g_i_*). Centre: the network graph in an equivalent adjacency matrix representation; green or red entry indicates the interaction is in agreement or disagreement, respectively, with the OA (e.g. *m*_14_ = −1 in disagreement with *a*_4_ = +1). Right: benefit/damage scores equal the sum of beneficial/damaging interactions *g_i_* is projecting onto or attracting from other genes (counting along row *i* for projection and column *i* for attraction; *g*_3_’s scores highlighted by dotted frames as an example). **(B)**, a knapsack optimization problem (KOP) instance; the challenge is to determine the objects to include in and exclude from the limited-capacity knapsack so as to maximize the overall total value while keeping the overall total weight of included items under the knapsack’s capacity threshold (5 lb). This KOP instance is reducible to the NEP instance in **A**; objects/values/weights/capacity correspond to the genes/benefits/damages in **A** (e.g. laptop in **B** corresponds to gene *g*_2_ in **A)**. Solving the question ‘which genes to conserve and which to delete’ in **A** translates into a solution to the KOP instance in B. NEP instance optimal solution, with a threshold ≤5 damaging interactions, is to conserve genes *g*_2_, *g*_4_, *g*_5_, *g*_6_ and delete *g*_1_, *g*_3_ *g*_7_, which respectively translate back to an optimal solution to the KOP instance: include in the knapsack the laptop, lunch box, pen and water bottle, and exclude from it the notebook, textbook, and candle. **(C)** and **(D)**, NEP can semantically be interpreted in the context of (C) regulation (which genes or interactions should be fine-tuned positively or negatively) or (D) evolution (which genes or interactions represent an asset or a liability to the system long-term). The formal definition of NEP and a generalized reduction of KOP to NEP are included in SI 3 and 4 respectively. See main text and SI 5 for interaction-targeting OA (as opposed to gene-targeting OA here in A).

The adjacency matrix *M* in Figure 2 (A), centre, is an equivalent representation of the network graph on the left. There is a non-zero entry *m_jk_* for each edge between genes *g_j_* and *g_k_*. A promotional or inhibitory interaction is represented in *M* as a +1 or −1 entry, respectively. A green- or red-coloured matrix entry *m_jk_* denotes whether the interaction is in agreement or disagreement with the OA on *g_k_*, i.e. *a_k_*. While *m_jk_* describes what the effect of *g_j_* on *g_k_* actually *is, a_k_* describes what that effect *should* ideally be.

For example, *g_1_* promotes *g*_2_ (*m*_12_ = +1) in agreement with the OA that *g*_2_ should be promoted (*a*_2_ = +1). On the other hand, *g*_1_ inhibits *g*_4_ (*m*_14_ = −1) in disagreement with the OA on *g*_4_ (*a*_4_ = +1). The benefit/damage (b/d) score of a gene *g_i_* is calculated as the sum of green/red interactions along row *i* (projection) and column *i* (attraction), as shown in the right tabular in Figure 2 (A). The benefit and damage scores of *g*_3_, and the corresponding row and column from which they have been summed up, are highlighted with dotted frame in Figure 2 (A). Notice that although the Oracle is neutral towards *g*_1_,*g*_5_ and *g*_6_, they do have b/d values depending on their interactions with other genes and whether those interactions are in agreement with the OA or not. Given the set of genes with non-zero benefit and damage score, the optimization problem (formally defined in SI 3) is:

*What subset of genes should be conserved and which should be mutated/deleted so as to maximize the total number of beneficial interactions network-wide, while minimizing (to a threshold) the total number of damaging interactions?*

## 4 The Computational Complexity of NEP

The computational complexity of NEP is equivalent to that of the well-known 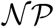-complete knapsack optimization problem (KOP). Figure 2 (B) shows a KOP instance with 7 items, each tagged with a certain value reflecting its utility, and some weight in pounds. The values and weights are indicated on items’ tags with green and red numbers respectively. The optimization question in KOP is how to pack the knapsack with as many items as possible such that the total value of items included in the knapsack is maximal while their total weight does not exceed the knapsack maximum capacity, which is 5 pounds in this KOP instance. Some items have negligible weight and some are useless, such as pen and candle in this example, respectively. Clearly the pen (candle) should be included in (excluded from) the knapsack regardless of what other objects’ fate, and as such need not be considered in the optimization search. Among the remaining objects, a search through the include/exclude combinations must be explored to determine the optimal solution.

We say that the KOP instance in Figure 2 (B) is reduced to the NEP instance in (A). Nodes *g*_1_,*g*_2_, …, *g*_7_ and their b/d scores in (A) respectively correspond to the items and value/weight scores of the KOP items (B). For example, KOP’s laptop corresponds to NEP’s gene *g*_2_. The optimal solution to the NEP instance in Figure 2 (A) would be to conserve *g*_2_,*g*_4_,*g*_5_ and *g*_6_ and to mutate or delete *g*_1_,*g*_3_ and *g*_7_, assuming the maximum threshold of tolerable damaging interactions to be 5 (corresponding to the 5 pounds knapsack capacity in (B)). This solutions respectively translate back to an optimal solution to the KOP instance in (B): include in the knapsack the laptop, lunch box, pen and water bottle, and exclude the notebook, textbook, and candle. Generally, any instance of KOP can easily (i.e. with polynomially bounded computational cost) be reduced into a corresponding NEP instance. A generalized formal KOP-to-NEP reduction is included in SI 4.

With small number of KOP items it is trivial to find out the optimal solution, but as the number of items increases the combinatorial search space grows exponentially fast. Whether there exists a polynomially-bounded algorithm for solving any arbitrary instances of an 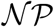-complete 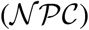 problems is the subject of the 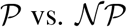 question, arguably the most important questions in computer science and mathematics today [18, 19, 20]. 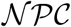 problems have defied all attempts aimed at finding algorithms that consume polynomially-bounded amount of computational resources for solving arbitrary instances. Hence, the increasingly accepted conjecture is that problems in this class will always require super-polynomial computational resources, and that this should be accepted as a universal law by the same token that repeatedly experimentally verified laws in physics are accepted as such [21]. In practice heuristics-based algorithms do exist and can find good approximation solutions (e.g. by exploiting structures in practically common instances) [22, 23]. In evolutionary context, however, the only ‘algorithm’ at hand is random variation non-random selection (RVnRS), and the NEP instance difficulty is measured relative to it.

## 5 Optimization in Biological Context

Biological systems do not employ sophisticated search algorithms to determine the optimal conserve or mutate/delete actions from one generation to the next, but rather proceed thru iterations of RVnRS [24]. However, the number of needed RVnRS iterations before the composition (nodes) and connectivity (edges) of a network has sufficiently been transformed away from a deleterious state depends directly on network topology. Particularly, the number of RVnRS iterations is exponential in the number of ambiguous genes (those having non-zero score for both benefit and damage). The more ambiguous a set of genes are, the less likely it is for the iterative RVnRS process to *alter just the right set of genes* such that the total number of beneficial/damaging interactions is are optimally maximized/minimized in as few evolutionary iterations as possible.

The 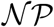-hardness of NEP in and of itself is a rather weak measure of computational difficulty as there are in practice heuristics-based approximation algorithms that can produce fairly satisfactory solutions that may not be too suboptimal relative to *the* optimal solution. However, the efficacy of RVnRS relies on how accumulating improvements with as little ‘backtracking’ through the search space as possible. The more unambiguously beneficial or damaging genes there are in a system that is under some evolutionary pressure to alter its MIN, the more effective RVnRS is at accumulating more beneficial, and cleansing more damaging, interactions.

A naive greedy strategy could be to impose the OA by conserving every gene ***g_i_*** where *a_i_* = +1, and mutating/deleting every *g_j_* where *a_j_* = −1. However, **conserving *g_i_*** can inadvertently conserve OA-contradicting interactions if *g_j_* happens to be a promoter or inhibitor of some *g_k_* where *a_k_* = −1 or *a_k_* = +1 respectively. And **deleting/mutating *g_j_*** can inadvertently disrupt OA-supporting interactions if *g_j_* happens to be a promoter or inhibitor of some *g_k_* where *a_k_* = +1 or *a_k_* = −1 respectively. In other words, a naive imposition of an OA is complicated by the reality of network connectivity. NEP optimization in biological evolutionary context is summarized in Table SI 4. Further details on the notions of gene ambiguity, search space size under RVnRS, and the role of mLmH topology at reducing it in Subsection 7.3 and 7.4.

## 6 The Semantics of NEP

The OAs described thus describes a rather blunt opinion about a gene by stating it should be promoted or inhibited. In reality a gene may ideally have both a promoter and an inhibitor co-expressed whereby, for example, the latter serves to tone down the potency of the former. A more fine-tuned OA would therefore be on individual interactions rather than genes. The only syntactic modifications to NEP definition in this case is that the OA is a matrix that is equal in dimensions to the interaction matrix *M*. This is in contrast to a gene-targeting OA which is represented as a |*G*|-long sequence where |*G*| is the total number of genes in the network (as was the case in the example of Figure 2 A). The complexity and instance difficulty of NEP remains the same whether the Oracle is stating its advice on genes or interactions, however (see SI 5 for further details).

An OA can further be interpreted semantically as describing short- or long-term evolutionary pressure. Short-term OA is specific to some regulatory state at a specific moment of the cell’s life cycle for example (Figure 2 (C)). An OA on some gene or interaction indicates an evolutionary pressure to fine-tune its regulation positively or negatively. Long-term OA on the other hand indicates a more existential evolutionary pressure for or a against a gene or interaction. In this case, the OA is interpreted as advising on whether a gene or interaction is rather advantageous or disadvantage to the organism’s survival generally (Figure 2 (D)).

## 7 Results

Here we summarize the results presented in subsequent subsections. We empirically study NEP instances obtained by generating hypothetical OAs on various MINs and compare that with instances on synthetic networks of various topologies. We assess instance difficulty assuming the working algorithm is RVnRS. The results show instances obtained from real MINs are invariably far easier to satisfy by virtue of the mLmH property, as real MINs have far less ambiguous genes per instance. The large number of low-degree leaf genes in MINs reduces instance size because such genes are certain (degree 1) or likely (degree 2, 3, .. with exponentially decreasing likelihood) to have all-beneficial (all-damaging)interactions and should therefore be conserved (mutated/deleted) regardless. Such unambiguous genes need not be considered in the computationally costly optimization search. For example, a leaf gene involved in only one interaction (i.e. it has degree 1) can either be totally beneficial or totally damaging under a given evolutionary pressure scenario, and as such there is no ambiguity as to whether it should be conserved or mutated/deleted. A gene of degree 2 has a 50% chance of being unambiguous: the two interactions it is engaged in are either both beneficial or both damaging (a degree-2 gene is engaged in two interactions. There are four possibilities of the state of the two interactions are (0,0), (0,1), (1,0) and (1,1) where 1 or 0 denote an interaction is beneficial or damaging respectively). In general, a gene of degree *d* has a 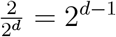 probability of being unambiguous.

Based on the fact that the larger a gene’s degree is the exponentially more likely it is to be ambiguous, the model predicts the expected number of genes of degree *d* in real MINs. The prediction formula is parameterized with a value proportional to the edge:node ratio of the MIN. All existing MINs are currently partial to different degrees of completion (i.e. not all interactions of all genes have been mapped out). Based on the predictability of the degree distribution of MINs from a computational intractability perspective, we conjecture that, once all genes and interactions have been accounted for, there will be a universal edge:node ratio of ~2 in all MINs regardless of organism or physiological context.

We also simulated the evolution of synthetic networks using an evolutionary algorithm which, in each generation, selects the top 10% of networks that present the easiest optimization task (mainly by having a large number of unambiguous nodes). These top networks breed the next generation of networks. The degree distribution of the fittest synthetic networks after ~hundreds - few thousands of generations of simulated evolution very closely match those of real MINs of equal size. The emergence of mLmH in synthetic networks is not sensitive to the starting conditions: whether networks are initially empty (and accumulate nodes/edges over the generations) or randomized (have the same number of nodes/edges as a corresponding real MIN, but edges are re-assigned over the generations as a form of mutation).

### 7.1 simulation of evolutionary pressure

NEP instances were generated on MINs (previously described in Figure 1, see also SI 2 for extensive details) as well as various synthetic networks. For each MIN, we computer-generated synthetic analog networks having the same number of node and/or edges. Figure 3 (A) shows the degree distribution of an example real network (*Drosophila melanogaster* (Fly) protein-protein interaction (PPI) network [25]). A no-leaves (NL) and no-hubs (NH) networks were generated by reassigning edges in PPI from leaves to nodes (for NL) or vice versa (for NH), so as to simulate the effect of depriving PPI of either property. Nodes in NL (NH) have a minimum (maximum) degree ≥ (≤) the average node degree of PPI (⌈~3.6⌈ =4). The redistribution of edges from leaves to hubs in NL results in some nodes having zero degree, which are eliminated, resulting in NL being a smaller (943 nodes), more dense network compared to PPI. NL and NH networks simulate two alternative topologies that biological networks could have evolved into if minimizing interactions per gene (NH) or total number of genes (NL) were the only driving forces in their evolution. We also applied the same simulation to a random (RN) analog of PPI network, whereby each edge in the latter is re-assigned to two randomly selected nodes (both edge direction and sign randomly assigned), and so nodes’ degrees cluster around the average degree in PPI network. Figure 3 (B)-(D) respectively show the degree distribution of analog (NL, NH, and RN) random networks for protein-protein, regulatory, and database-sourced networks (the degree distribution of corresponding real networks were shown in Figure 1). 1-5K NEP instances are generated for each network by calculating *B* and *D* values against a randomly generated OA on all nodes (for each node *n_i_*,*a_i_* ≠ 0). Each instance is solved to optimality under a given tolerance threshold, expressed as the % of damaging edges to be tolerated. Details of the algorithmic workflow of the simulation, and the choice of sampling threshold, is provided in SI 6, 7, respectively.

**Figure 3:**
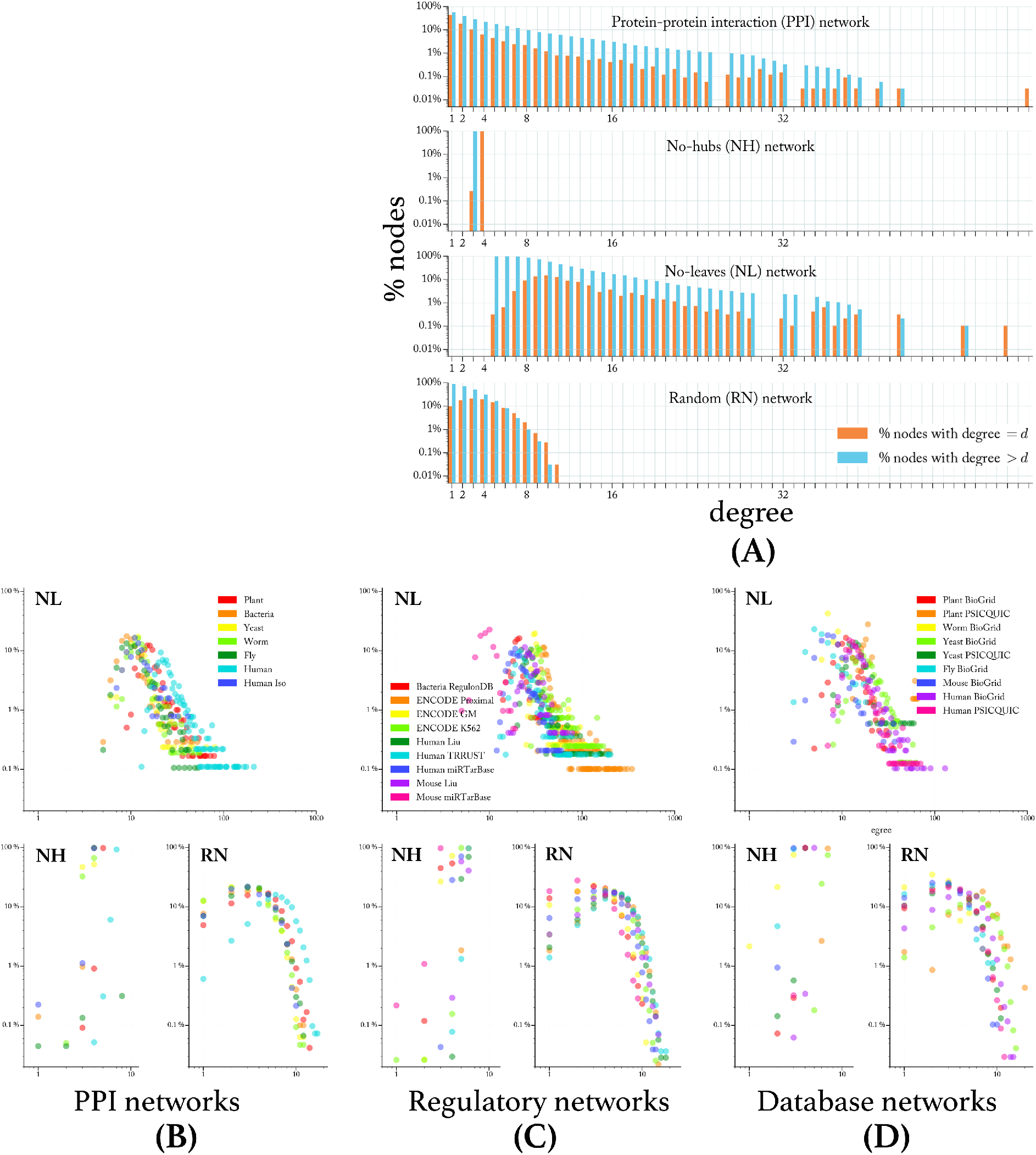
Case study networks. **(A)** Degree distribution of an example MIN (Fly protein-protein interaction (PPI) network) and its corresponding synthetic analogs; orange (blue) bars represent the % of nodes with degree = *d* (> *d*). NL (NH) networks have a minimum (maximum) degree > (≤) the average degree in the PPI network (~ = 4). PPI nodes have majority low-degree nodes (~78% with degree ≤ 4); degrees in RN cluster around the average degree due to re-assignment of each edge in PPI to two nodes selected uniformly randomly. **(B-D)** The degree distribution of synthetic analog networks corresponding to each real MIN from the protein-protein (B), regulatory (C), and database-sourced (D) categories as shown in Figure 1.

### 7.2 benefit-damage correlation

The correlation between benefit and damage scores is used to assess difficulty of NEP instances analogously to values-weights correlation in KOP (see NEP-to-KOP reverse-reduction in SI 4.3). Figure 4 (A) shows the correlation plot of classical test instances [26]. It has previously been shown that the more correlated the values and weights in a knapsack instance are the more difficult the instance is [27]. Strong value-weight correlation increases the ambiguity as to which items to add/remove from the knapsack (or in NEP context, which genes to conserve/delete from the network). Figure 4 (B) shows the average frequency of a (benefit, damage) pair ((*b*, *d*) hereafter) over 1-5K NEP instances for each network. The reduced ambiguity in instances from real PPI networks results by virtue of the large number of genes that are certainly (degree 1) or likely (degree 2, 3, 4 .. with likelihoods 50, 12.5, 0.125 .. %, respectively) to be unambiguous: either totally advantageous (*b* ≠ 0, *d* = 0) or totally disadvantageous (*b* = 0, *d* ≠ 0). In Fly PPI network for example, ~50% of all (*b*, *d*) pairs are unambiguous, with 1:0 and 0:1 pairs (resulting from degree-1 leaf genes) alone representing ~43% of those pairs (large brown dots). In contrast, ~15.6, ~3.1, and ~28% of (*b*, *d*) pairs in Fly network’s NH, NL, and RN analogs are unambiguous. The role of leaves in decreasing (*b*, *d*) correlation is especially highlighted in contrast to leaf-deprived NL network which exhibits the strongest correlation around its higher mean degree (analysis of degree-to-ambiguity proportional relation is detailed in “Prediction of degree distribution” section). The symmetry along the diagonal in Figure 4 (B) is expected given that there is an equal probability of 50% that an interaction is beneficial or damaging under a random Oracle advice on the gene that interaction is targeting.

**Figure 4:**
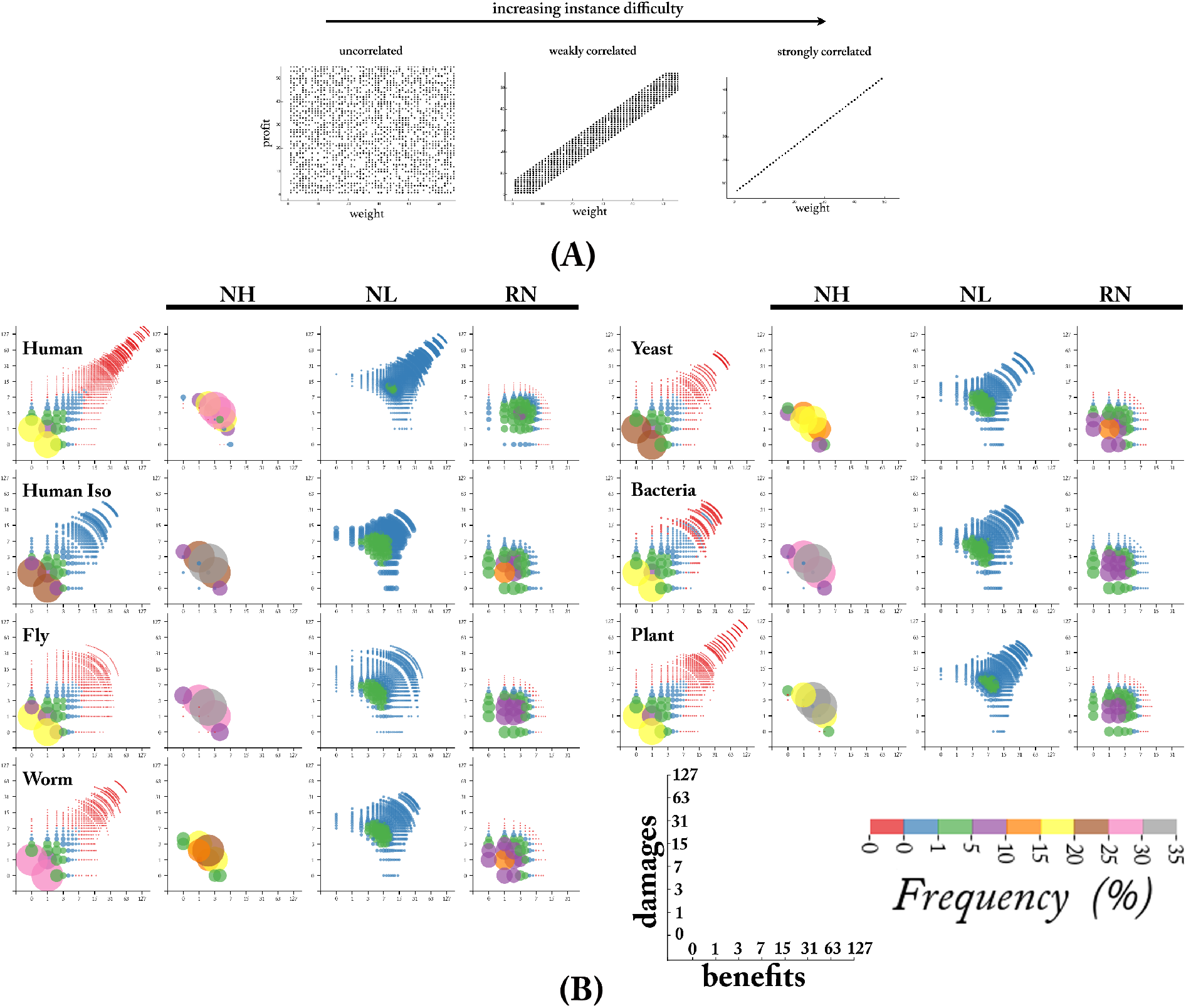
Benefit-damage correlation as an instance difficulty measure. **(A)** Value-weight correlation in classical knapsack instances; the stronger the correlation the computationally harder the instances [27]. **(B)** Benefit-damage correlation in NEP instances from PPI networks and corresponding synthetic analogs. The % of genes having a given (benefit, damage) score, (*b*, *d*), in an NEP instance (averaged over 1-5K instances). Size and colour of each dot reflects frequency of that (*b*, *d*) pair (see bottom-right legend). ~50% of all (*b*, *d*) pairs in Fly PPI network for example are unambiguous (*b* = 0,*d* ≠ 0 or *b* ≠ 0,*d* = 0) largely due to leaf genes of degree 1 (alone contributing on average ~43%, large yellow dots in Fly subplot). In contrast, the fraction of unambiguous (*b*, *d*) pairs in NH, NL, and RN are ~15.6, ~3.1, and ~28%, respectively, as their most frequent (*b*, *d*) pairs cluster around dominant degrees (a (b,d) pair is contributed by nodes of degree *d*=*b*+*d*). Leaf-deprived NL network manifests the strongest (*b*, *d*) correlation given the range of ambiguity that most of its nodes (clustered they are around NL’s relatively higher mean of ~12) can assume. The symmetry along the diagonal is expected given that there is an equal probability of 50% that an interaction is beneficial or damaging under a random Oracle advice on the gene that interaction is targeting.

The signed Fly PPI network distinctly shows more sporadic benefit-damage correlation (red dots in Fly dot plot in Figure 4) in higher-degree nodes, which results from the asymmetry of the number of promotional (67.5 %) and inhibitory (32.5 %) interactions. Under random OA, such asymmetry results in higher likelihood of disparate benefit and damage values. The asymmetry of signs in Fly PPI network is consistent with a recent report that showed a similar promotional/inhibitory interaction distribution in yeast [28]. The benefit-damage correlation for regulatory and DB-sourced networks manifest the same property (see SI 8).

### 7.3 effective instance size

Unambiguous genes can *a priori* be deemed advantageous or disadvantageous and therefore should be conserved or mutated/deleted, respectively. As such they need not be part of the optimization search because the optimal evolutionary outcome as to whether to conserve or alter them is independent of that of other genes in the network. Effective instance size (EIS) is the fraction of genes in an NEP instance that are ambiguous (*b* ≠ 0 and *b* ≠ 0). The smaller the |*b* − *d*| value the more ambiguous a gene is. Figure 5 (A), left (pie charts) show the fraction of genes that on average (over 1-5K instances) falls under a certain *b*:*d* ratio slice where 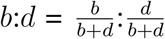 for an example network (Fly PPI). NEP instances in PPI have ~42% EIS (*b*:*d* = 90:10, 80:20, .., 10:90%), compared to ~84, ~100, and ~72 % for NH, NL and RN networks respectively. Compared to NH and NL, EIS in RN is smaller to the extend that it has more leaf nodes (particularly degree 1-3) which relatively increase the size of its unambiguous slices (*b*:*d* 100:0 or 0:100%).

**Figure 5:**
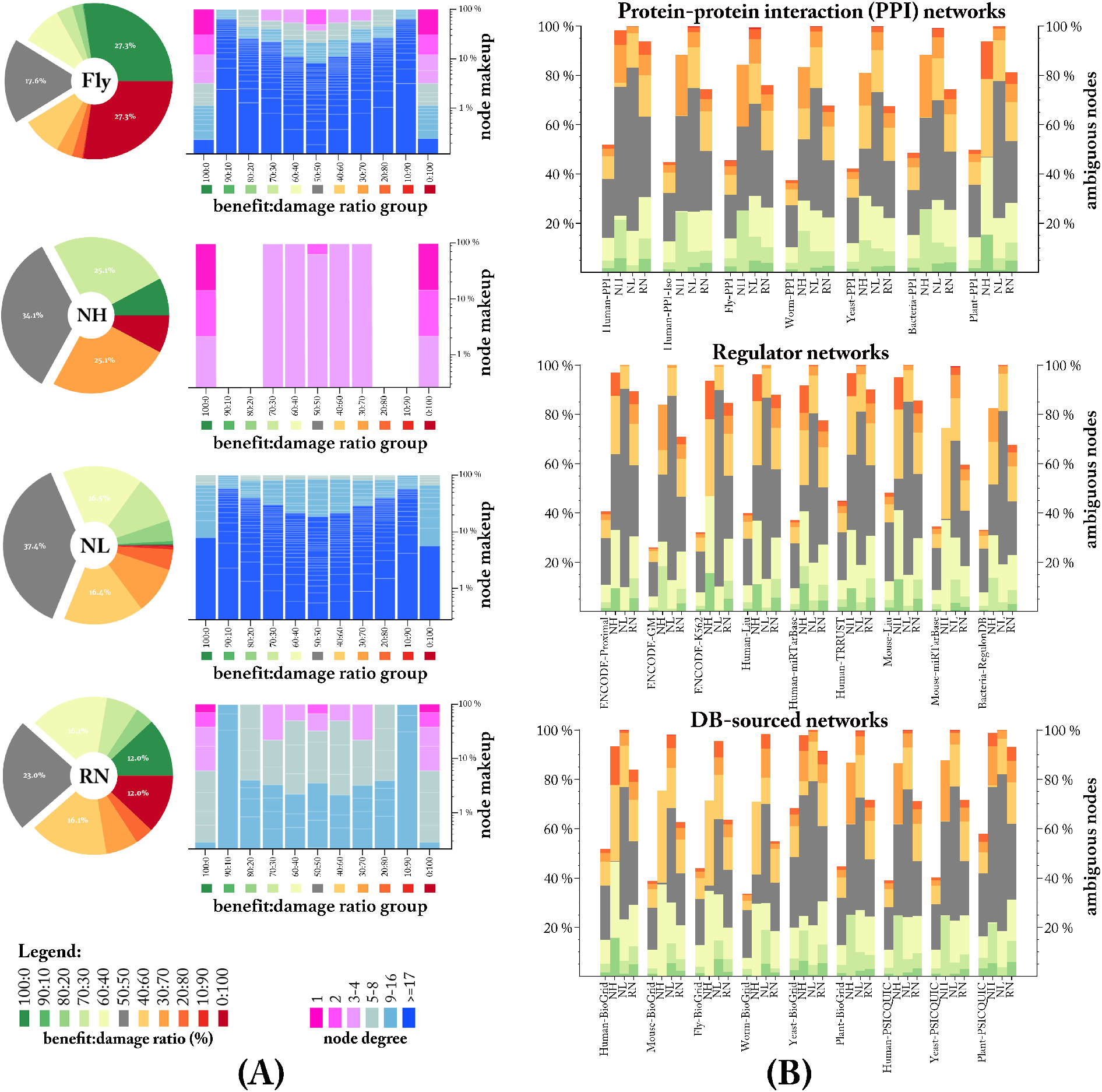
Effective instance size (EIS). **(A)** left pie charts: fraction of nodes in NEP instances having a certain benefit:damage (*b*:*d*) ratio (bottom legend) in the Fly PPI network and its synthetic analogs. The subset of nodes that need to be optimized over (those in all but 100:0, 0:100 slices) is significantly smaller in leaf-rich PPI (Fly) instances. Virtually all nodes in leaf-deprived NL network are ambiguous (~0% nodes under 100:0 or 0:100 *b*:*d* ratios); right bar charts: break down of genes contributing to each *b*:*d* ratio slice in the corresponding pie chart, broken by gene degree (bottom legend); leaves dominate 100:0 and 0:100 slices in PPI. Because all nodes in NH network have degrees ≤ 4, no *b*:*d* ratio of any gene can fall in the 90:10/10:90 or 80:20/20:80 slices. **(B)** EIS for all networks; bar height represents the ambiguous pie slices in (A); each bar group corresponds to a MIN along with its corresponding NH, NL and RN analogs. In all networks, EIS is significantly smaller in real MINs compared to synthetic analogs. RN networks have more leaves (by sheer random re-assignment of edges in their creation) and therefore have smaller EIS compared to NL and NH.

The constituent genes in each pie slice are shown in Figure 5 (A), right bar charts, for each network, broken down by degree range (bottom legend). Since the likelihood of a gene’s ambiguity is inversely (and exponentially, see later discussion in the next section) proportional to its degree, leaf genes (degree ≤ 4) in PPI dominate the unambiguous 100:0 or 0:100% *b*:*d* ratio groups. With virtually all nodes in NH network having a degree 4 (NH bar chart in Figure 3 (A)), no *b*:*d* ratio of any node can fall in certain *b*:*d* ratio groups (more specifically, none of the possible (b,d) pairs 4:0, 3:1, 2:2, 1:3 … 0:4 falls into any of the *b*:*d* ratios 90:10, 80:20, 10:90, or 20:80 %). Figure 5 (B) shows the EIS for all networks, where the bar height represents the size of ambiguous pie slices shown in (A) excluding the 100:0 and 0:100 slices. Each bar group corresponds to a network, with the first bar being the real network and the 2nd, 3rd and 4th (left to right) corresponding to the network’s NH, NL and RN analogs respectively. EIS is significantly smaller in real networks as compared to synthetic analogs regardless of physiological context or the source of the network. EIS in RN analogs is comparatively smaller in networks having smaller edge:node (*e*2*n*) ratio since such networks are more likely to have leaf nodes after random re-assignment of edges (e.g. Human and Yeast PPI networks have 1.45 and 3.24 *e*2*n* ratios respectively).

### 7.4 search space size

The total search space of a given NEP instance is defined as *S_t_* = 2^*O*(*n*)^ where *n* is the number of relevant genes in the instance (those with *b* ≠ 0 and *d* ≠ 0). The base 2 denotes the two general evolutionary outcome on a relevant gene *g_i_*: its state will have been (1) altered or (2) unaltered after the next round of RVnRS. The alteration to a gene (for example through mutation) can be (dis-)advantageous relative to the current NEP instance. An advantageous change includes for example the deletion of a unambiguously damaging gene. The state of an ambiguous gene can further be affected by the state changes of its interacting partners. For example, in the NEP instance of Figure 2 (B), a mutation to gene *g*_2_ which causes it to lose its positive interactions with say, *g*_1_ and *g*_5_ would not only change its benefit/damage (*b*, *d*) scores from (4, 2) to (2, 2) but it would also result in the (*b*, *d*) scores of *g_1_* and *g*_5_ to change even if they themselves did not undergo any alteration. *g_1_* and *g*_5_ lose 1 benefit as a result of them no longer positively interacting with *g*_2_ due to the latter’s aforementioned mutation.

Clearly the more interconnected a gene is the more its state alteration, and the alteration of its interacting partners, changes the optimal evolutionary target for such a gene from one generation to the next (i.e. whether it should ideally be conserved or mutated/deleted given the current evolutionary pressure described by the OA). Effective search space is defined as *S_e_* = 2^*O*(*m*)^ where 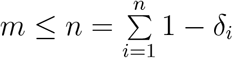 and 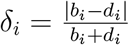 is the ambiguity measure of *g_i_* under the current NEP instance. On the one extreme where ∀*g_i_*, *b_i_* = *d_i_* ≠ 0, then *m* = *n* and therefore *S_e_* = *S_t_*. On the other extreme where all genes are unambiguous, ∀*g_i_*, *b_i_* = 0|*d_i_* = 0, then *S_e_* = 0. An unambiguously beneficial or damaging gene *g_j_* (*d_j_* = 0 or *b_j_* = 0, respectively) does not increase *m* regardless of the current evolutionary pressure scenario because *δ_j_* = 1 (gene ambiguity vs. degree is discussed in details in the subsequent section “Prediction of degree distribution”). Figure 6 shows *S_t_* (grey) and *S_e_* (blue) for all networks. Note that the *y*—axis values are logarithmic in base 2, and hence linear differences between bar heights imply exponential differences in *S_t_* or *S_e_*. The smaller but denser NL networks show smaller *S_t_* but suffer from exponentially higher *S_e_*:*S_t_* ratio compared to other networks. This implies that, while the search space is smaller, the optimal solution can change radically after each RVnRS given how deeply interconnected the genes are in such a network. Biological networks show exponentially smaller *S_e_*:*S_t_* ratio especially for larger more complete (e.g. ENCODE and Human PPI connectome) and/or curated (e.g. TRRUST) networks.

**Figure 6:**
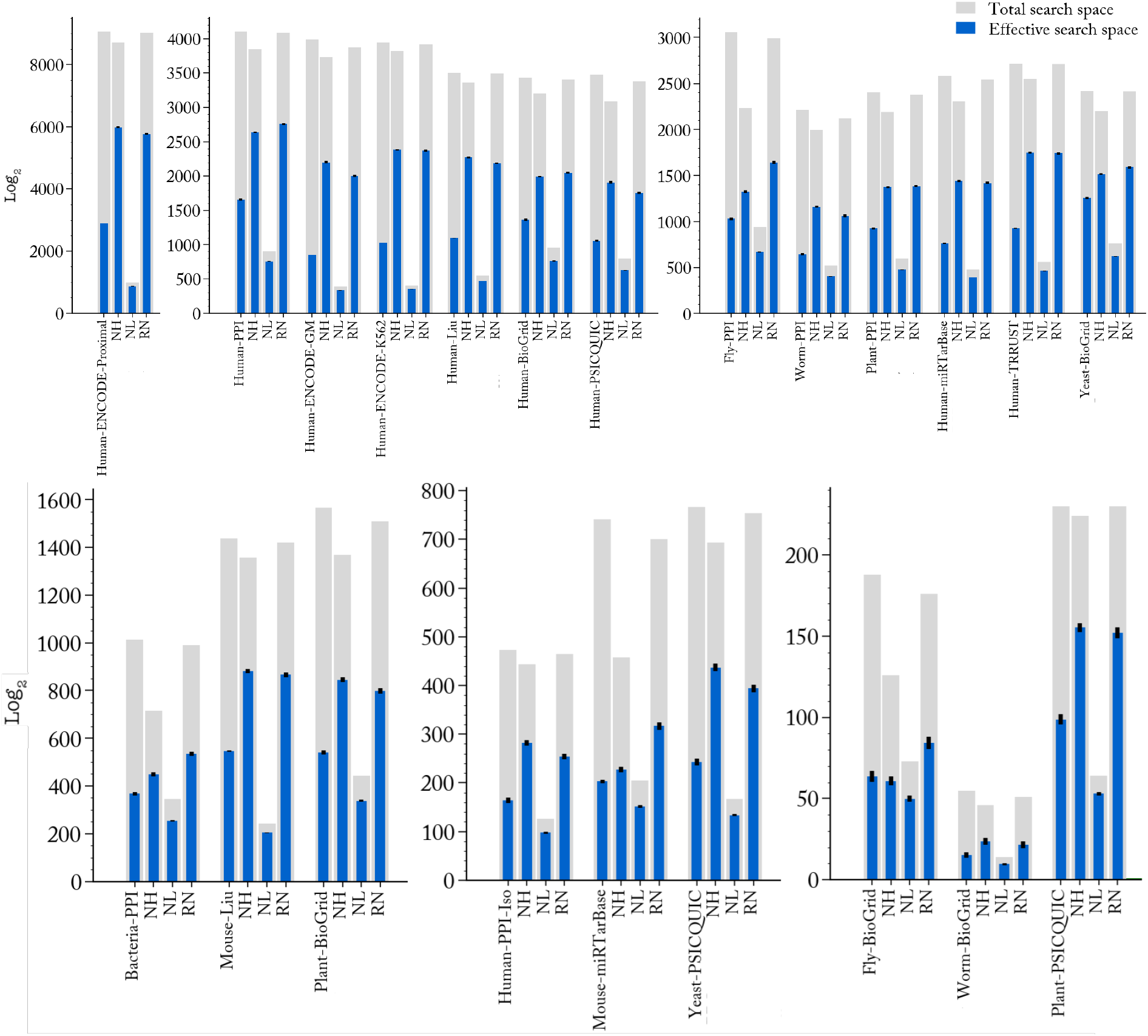
Search space size. The total search space (grey) of a given NEP instance is quantified as *S_t_* = 2^*O*(*n*)^ where *n* is the number of genes in the instance with either benefit score *b* > 0 or damage score *d* > 0. Effective search space (blue) is defined as *S_e_* = 2^*O*(*m*)^ where 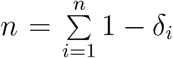 and 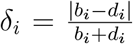 is the ambiguity measure of *g_i_* under the current NEP instance (*b_i_*, *d_i_* are the benefit, damage scores of gene *g_i_*). Networks are grouped by size for better *y*-axis readability. Note that the *y*—axis values are logarithmic in base 2, and hence linear differences between bar heights imply exponential differences in *S_t_* or *S_e_*. Smaller but denser NL networks have smaller *S_t_* relative to other networks but suffer from having extremely high *S_e_*:*S_t_* ratio due to the high ambiguity of its nodes under NEP instances. Biological networks have exponentially smaller *S_e_*:*S_t_* ratio relative to corresponding NL, NH, and RN random analogs by virtue their having a higher number of leave nodes, particularly genes of degree 1-3, which are more likely to be unambiguously beneficial or damaging under a given NEP instance, and as such have relatively higher *δ* values (= smaller *m* exponent overall).

### 7.5 gene neutrality

To test the affect of having a certain fraction of the genes as neutral under a given evolutionary pressure, NEP instances were generating assuming 0, 25, 50, and 75% of genes are neutral (the Oracle has no opinion about them). A gene towards which the Oracle is indifferent may nonetheless end up having a non-zero benefit and/or damage value depending on its interaction profile with other genes on which the Oracle has an opinion. Subplots from top-left (0%) to bottom-right (75%) in Figure 7 (A) demonstrate the resulting effective instance size (EIS) as pressure decreases (i.e. as the number of neutral genes increases). The bar height represents the effective instance size (EIS), i.e. the fraction of genes that are ambiguous (in that they are engaged in both beneficial and damaging interactions in a given NEP instance). EIS shown here is the average over 1-5K instances. As the pressure increases (bottom to top), the supremacy of real MINs compared to synthetic analogs (no-hubs (NH), no-leaves(NL) and random (RN) networks) is more pronounced. Leaf-deprived NL synthetic analogs show the largest EIS across all settings, while NH and RN networks show decreasing EIS to the extent that they both have more smaller-degree leaf nodes. The constituent nodes in each *b*:*d* bar segments is shown in Figure 7 (B) for the Fly network and its corresponding random analog. As the pressure decreases from 0% to 75%, the constituent nodes contributing to a given *b*:*d* ratio range remains largely unchanged as the nodes towards which the Oracle is not indifferent are selected randomly and as such the relative frequency of nodes of a certain degree remains of the same proportion under different neutrality assumptions.

**Figure 7:**
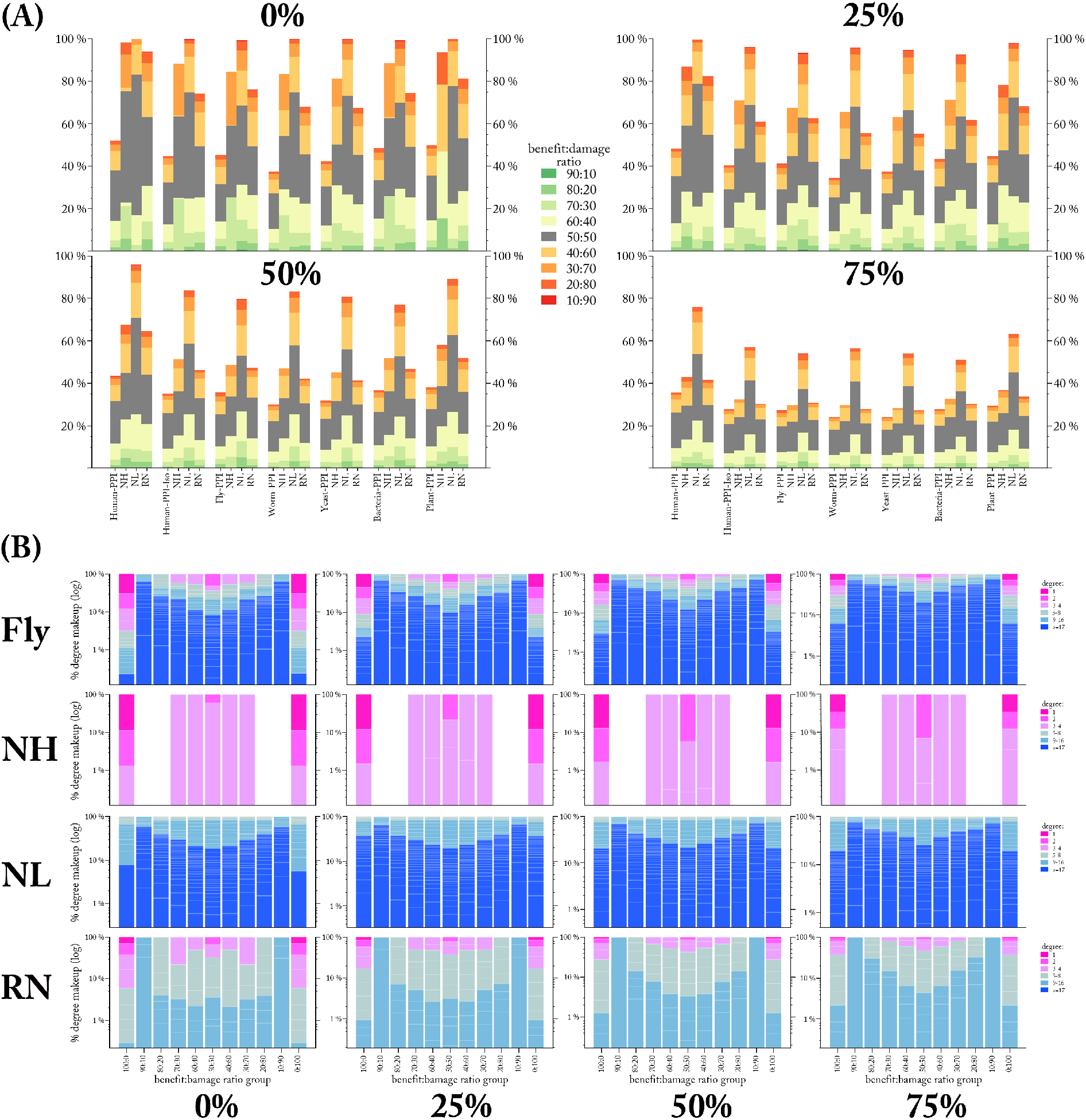
Neutrality of genes under evolutionary pressure. **(A)** NEP instances are generated with 0, 25, 50, and 75% of genes being neutral (the Oracle has no opinion on them). A neutral gene may nonetheless end up having a non-zero benefit and/or damage score in an NEP instance depending on its interactivity with non-neutral genes. The bar height represents the average EIS (see Figure 5). As the pressure increases (i.e. as the percentage of neutral genes decreases), the supremacy of real MINs compared to random analogs (no-hubs (NH), no-leaves(NL) and random (RN) networks) is more pronounced. **(B)** A zoomed-in view to the constituent nodes in each *b*:*d* ratio range for the Fly simulations. In the real MIN (Fly), as the pressure increases from 75% to 0% of genes being neutral, the utility of leaf genes becomes more pronounced as they dominate the 100:0 and 0:100 slices.

### 7.6 prediction of degree distribution

We considered whether computational intractability alone can predict the degree distribution of a biological network. More precisely, we considered whether the likelihood of a gene of degree *d* (“degree-*d* gene” hereafter) to be totally advantageous or disadvantageous (belonging to green or red pie slices in Figure 5 (A), respectively), which is exponentially inversely proportional to its degree, can predict the expected number of degree*-d* genes in a biological network. For a degree-2 gene *g_i_*, for example, there are 2^2^ = 4 potential states of benefits/damages that *g_i_* can assume under a given OA: 00, 01, 10, or 11 where 0 or 1 signify the edge (interaction) as being beneficial or damaging, respectively. States 01 or 10 are “ambiguous”: *g_i_* must be part of the overall optimization search to determine whether to conserve or delete it. In general, the number of ambiguous states for degree*-d* gene is 2^*d*^ − 2, albeit not all of equal ambiguity: while the 1000010 and 1111000 states of a degree-7 gene are both ambiguous, the former is significantly less so. Let *k* correspond to the number of 1’s in a gene’s given state (equivalently, its benefit score in a given NEP instance). We refer to a given state of a degree-*d* gene as *k-ambiguous* (*k-amb* hereafter), 0 ≤ *k* ≤ *d*, if it has *k* 1’s. For example, 0111 and 1000 are 3-*amb* and 1-*amb* states of a degree-4 gene. As *k* → *d* (or *k* → 0) the ambiguity whether to conserve (or delete) decreases, while as 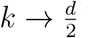 both the ambiguity and (exponentially) number of states increases. For a degree-20 gene for example, there are 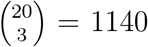 3-*amb* states compared to 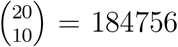 10-*amb* states. Assuming an equal probability 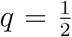 for an edge to be beneficial or damaging, the likelihood of *k-amb* state for degree-*d* gene is given by the expected number of *k* successes in *d* Bernoulli trials:

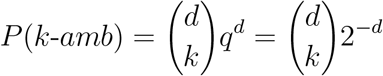

We define the expected frequency of degree-*d* genes as:

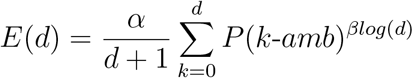

where constants *α*, *β* ∈ ℝ^+^ are proportional to node:edge (*n*2*e*), edge:node (*e*2*n*) ratios, respectively. Figure 8 (A) shows the actual (pink) and predicted (green) degree frequency in Fly PPI network at *α*, *β* = 0.43,1.9, respectively. Prediction is further applied to all other networks with accuracy (defined as 100 - ∑|*predicted*(*d*) − *actual*(*d*)|) being >= 84% as shown in Figure 8 (B). Individual prediction plots for all networks is included in SI 9. An accurate account of all interactions currently is affected by the experimental bias against interactions involving lesser known genes (an inherent problem to small-scale studies [1]). An accurate account of all genes is not only limited by experimental coverage but also by alternatively-spliced isoforms of the same gene (which can have distinct interaction profiles [2]) being treated as a single gene, hence a gene’s degree may be inflated in experiments where isoforms are not distinguished. For example, nodes in HumanIso network can be different isoforms of the same gene [2]. The (*α*, *β*) values versus (*n*2*e*, *e*2*n*) ratios in a partial MIN should indicate how well its coverage and resolution compares to other standard high-quality MINs. Figure 8 (C) shows (*α*, *β*) versus (*n*2*e*, *e*2*n*) ratios of networks in (B) (all of which are currently partial, see SI 2). The average of (*α* vs. n2e), (*β* vs. *e*2*n*) are (0.43±0.063 vs. 0.526±0.133), (1.96±0.324 vs. 2.06±0.634) respectively. We conjecture that there is ultimately (as experimental coverage and resolution of high-throughput interaction-detecting experiments increases) a universal *e*2*n* ratio of ~2 in all MINs.

**Figure 8:**
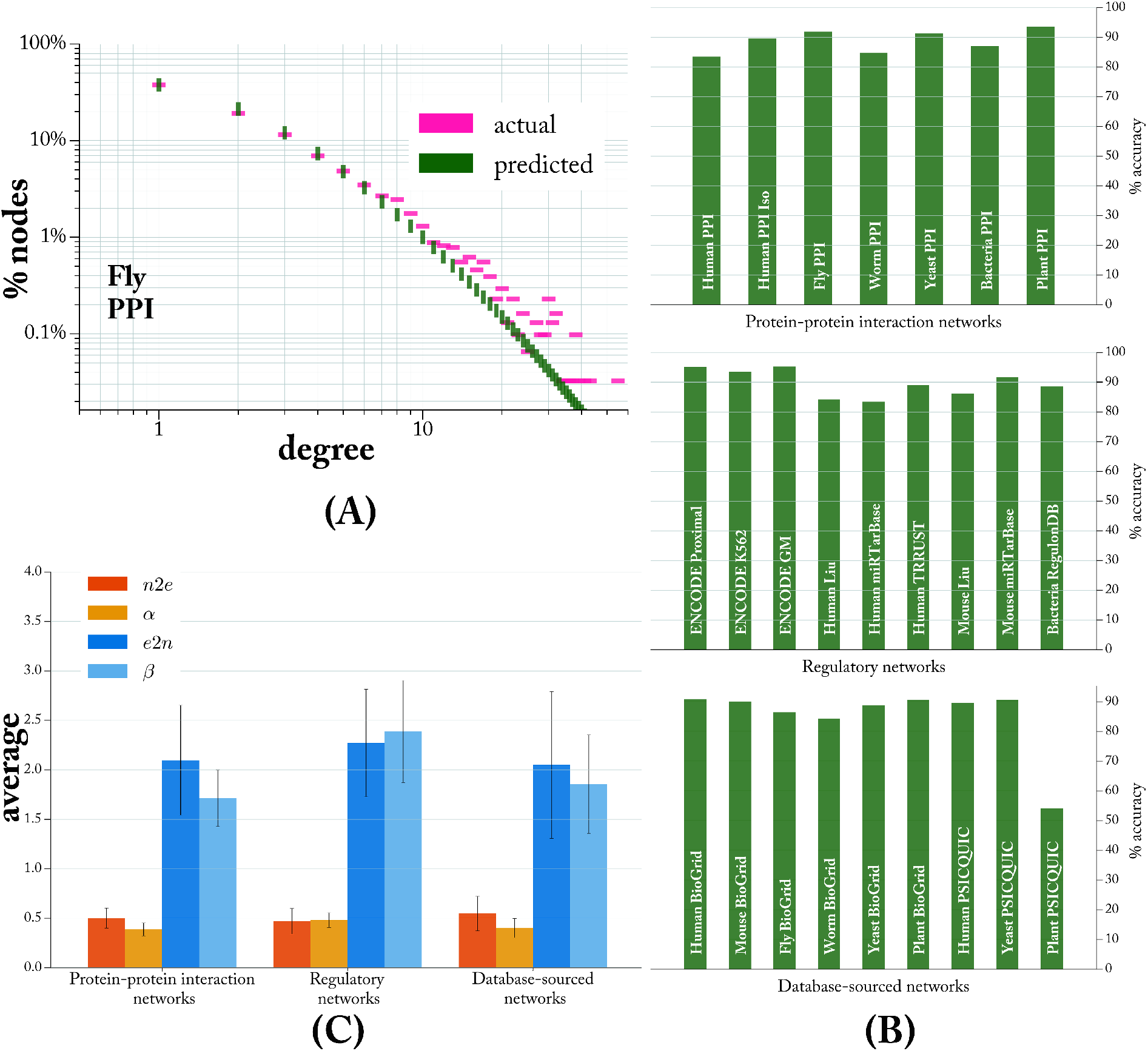
Computational intractability as a predictive tool of degree distribution. **(A)** The percentage of nodes having a degree *d* in Fly PPI network; the fraction of degree-*d* nodes is inversely proportional to the potential optimization ambiguity that a degree*-d* node adds to instances of NEP (see text). **(B)** Accuracy of predicting the degree distribution of PPI (top), regulatory (middle) and DB-sourced (bottom) networks (degree distribution plots of all networks is included in SI 9). Accuracy = 100 - Σ |*predicted*(*d*) − *actual*(*d*)| over each degree *d* in the network (*predicted*(*d*) = *E*(*d*) (see text) and *actual*(*d*)=the fraction of genes having degree *d*). **(C)** Proportionality of *α*, *β* to edge:node (*e*2*n*) and node:edge (*n*2*e*) ratios (*n*2*e* = *e*2*n*^1^), respectively, in the prediction formula *E*(*d*). The average±SD of (*α* vs. *n*2*e*), (*β* vs. *e*2*n*) are (0.43±0.063 vs. 0.526±0.133), (1.96±0.324 vs. 2.06±0.634) respectively.

### 7.7 simulated evolution

We conducted simulated network evolution and adaptation using an evolutionary algorithm that selects for networks that best minimize the difficulty of NEP instances generated against them. The simulation aims to assess the potency of NEP as a selection pressure that shapes network topology, rather than to mirror the details of recombination and mutation processes in biological systems. We conducted two sets of simulations depending on the starting conditions of synthetic networks. In the first, we begin with empty networks that grow over the generation by accumulating more nodes and edges. In the second, we begin by synthetic networks that have the same size as a corresponding real MIN, which is kept unchanged from one generation to the next except that an edges is re-assigned randomly in each generation as a form of mutation. We refer to the first and second sets of simulations as ‘evolution’ and ‘adaptation’ simulations respectively. In both experiments, the idea is to examine the degree distribution of synthetic networks after iterations of random mutation non-random selection, and compare that to the degree distribution of biological networks of equal size.

In simulated evolution, networks at the beginning of the simulation are near empty but are periodically assigned more new nodes and edges, in addition to being mutated by randomly reassigning an existing edge to two randomly selected nodes. The simulation begins with a population of 60 near empty synthetic networks (8 nodes and 8r randomly assigned edges, where *r* is the edge:node (*e*2*n*) ratio to be maintained throughout the simulation). An NEP instance against each network in the population is obtained by generating a random interaction-targeting OA, resulting in some interactions in the network being beneficial and others damaging. Nodes’ benefit/damage scores are calculated (as illustrated previously in Figure 2, and further detailed in SI 5). Assuming a tolerance threshold of 5% of total damaging interactions network-wide, instances are solved to optimality. (detailed further in SI 6).

The fitness of each synthetic network is based on two values averaged over the 60 instances: 1) effective instance size (EIS) as described in Section 7.3 and 2) the effective gained benefits (EGB) which measures the total number of beneficial interactions than can be obtained with the least number of nodes having to be conserved/deleted in an optimal NEP instances solution (see SI 10 for details). The top 6 fittest networks (10% of the population) that best minimize EIS and maximize EGB survive, while the remaining die. The surviving 6 networks are each mutated (add-node, add-edge, and re-assign edge) and 10 exact replicas are bred from each for the next round of mutate-and-select, resulting in a population of 60 networks once again. Another round starts by generating 100 NEP instances against each of the 60 networks in the population, and so on. The simulation is terminated when the number of nodes reaches that of a corresponding real MINs. Throughout the simulation, the add-edge mutation is conducted only if the edge-node ratio (*e*2*n*) of the synthetic network does not exceed that of the corresponding real MIN.

Figure 9 illustrates a sample of evolved synthetic networks and the corresponding MINs that have the same *e*2*n* ratio (results for all other networks are shown in SI 10). The degree distributions of simulated networks (blue in Figure 9) closely match their corresponding MINs (pink). In simulated adaptations, the starting synthetic networks have the same size as the corresponding MIN, and no add-node or add-edge mutations are conducted (i.e. synthetic networks don’t grow in size from one generation to the next). Only reassign-edge mutation is conducted in each generation. All other aspects of the mutation/selection process is the same as in the aforementioned simulated evolution experiments. mLmH property still emerges (see plots in SI 11). These results show the sufficiency of NEP-based evolutionary pressure to mold the network to mLmH topology within a number of generations that is extremely small (proportional to the number of genes in the target real MIN, i.e. ~thousands, see SI 2 for network sizes) relative to real evolutionary time. These results also confirm findings from previous simulation and adaptation experiments on a smaller set of networks reported in [29].

**Figure 9:**
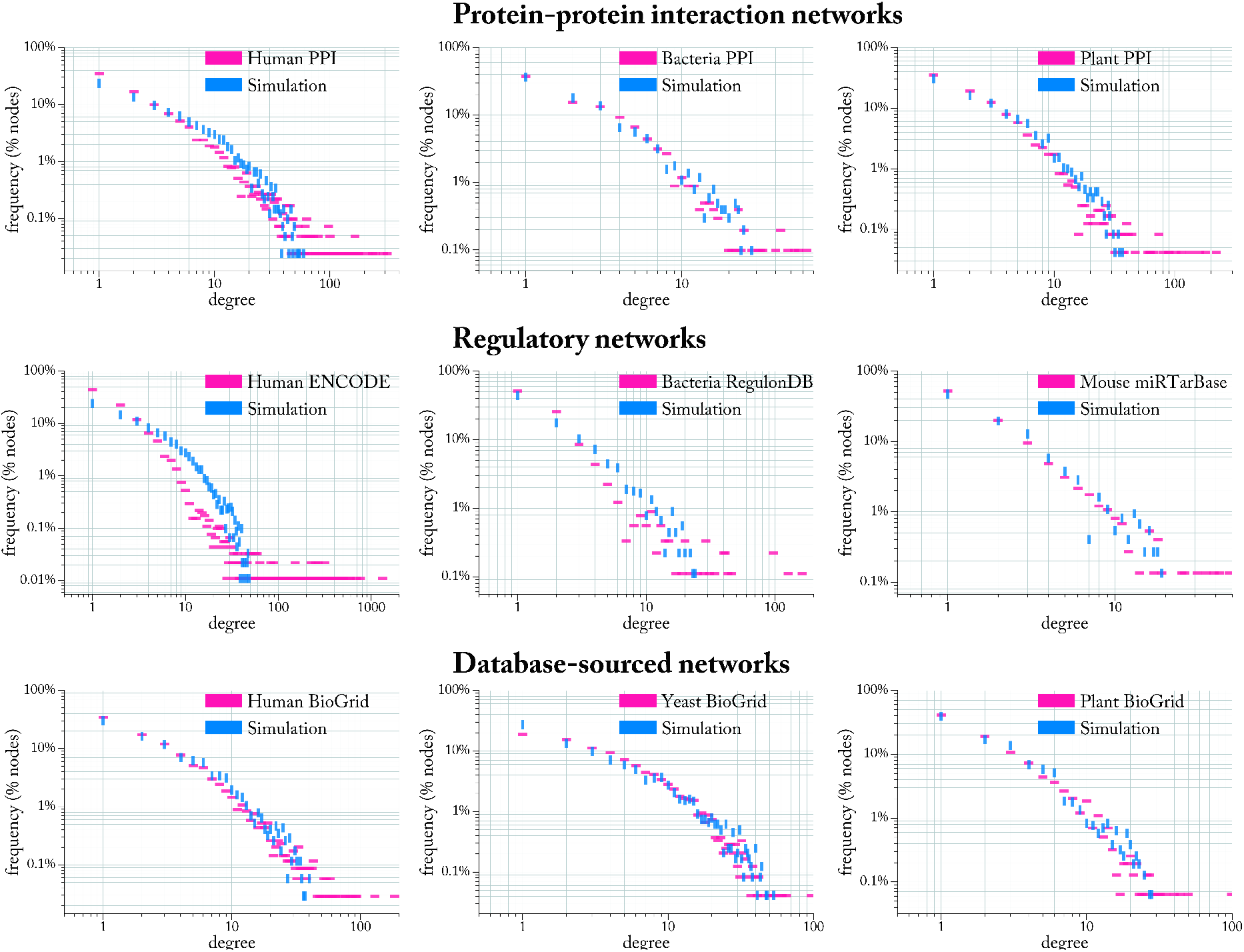
Evolvability under NEP selection pressure. Networks start empty and periodically undergo reassign-edge, add-node, add-edge mutations. An evolving network grows by adding one node, and one or more edges while maintaining an edge:node ratio equalling that of the corresponding real MIN. The simulation terminates when networks reach the same size (number of nodes) as that of the corresponding real MIN. The final degree distribution of the fittest network is illustrated (vertical blue dashes) against that of the corresponding MIN (horizontal pink dashes). Networks in row 1, 2 and 3 are protein-protein, regulatory, and database-sourced, respectively. The complete results for all other networks is included in SI 10.

## 8 Concluding Remarks

### contribution

The power of random variation and non-random selection to produce fine-tuned ‘hardware’ has been extensively documented at various biomechanical levels. Classic examples include allometric scaling and anatomical adaptations (organism), width and material of blood vessels (organ), compartmentalization of sub-cellular components (cellular) and fast-folding and aggressively-functioning enzymes (molecular). Recent reports [30, 31] have highlighted evidence for ‘software’ optimization in biological systems from a computational complexity perspective, proposing justification for the evolutionary advantage for the role of sex for example [32]. Such investigations into evolutionary biology through the lens of computational complexity have hitherto been too high-abstracted. Nonetheless, they do demonstrate that computational-complexity perspective has potential to be the much-needed theoretical framework that can guide the process of turning massive volumes of biological datasets into actionable knowledge [33].

There is currently a gap between the ever increasing scale and quality of molecular interaction networks (MINs) and the oretical understanding of the origin of their architectural properties. It has long been debated whether properties such as the majority-leaves minority-hubs (mLmH) is (non-)adaptive. Here we showed how computational intractability can provide sufficient and necessary conditions for the emergence of mLmH as an adaptation to circumvent computational intractability. The model predicts and evolves the mLmH based on the fact as a gene’s degree increases linearly, its optimization “ambiguity” potential increases exponentially and hence the more computationally expensive is the task of adaptively re-wiring the network away from a deleterious and into an advantageous state. We conjectured based on predictability results that there is ultimately a universal edge:node ratio of ~2 in all MINs. Given the increasing pace at which the coverage [34, 1] and resolution [35, 2] of high-throughput interaction-detecting experimental procedures is being reported, we expect the validity of such conjecture to be tested within a few years.

### sufficiency and necessity

We described the need for biological networks to change their node (gene) and edge (interaction) composition in response to some evolutionary pressure as a computational optimization problem which we showed to be 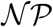-hard. By itself, the 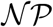-hardness of a problem is a rather weak measure of computational difficulty. Instances dealt with in practice may possess exploitable properties and as such their optimal solutions could potentially be routinely found without incurring exponential computational resource expenditure [22]. There could also exist good approximation algorithms that can solve to various degrees of optimality while consuming sub-exponential computational resources [23]. However, in the context of biological evolution, the running algorithm is random-variation and non-random selection (RVnRS) and the instance difficulty of the network re-wiring problem must be assessed relative to it. Our results show that the search space size for the described network re-wiring problem would be exponentially orders of magnitude larger were the topology of biological networks to deviate significantly from mLmH.

While the model sufficiently predicts and generates the mLmH topology, it also provides a necessary condition for the emergence of mLmH as an adaptation around inescapable computational intractability considering that (1) the computational cost of the network rewiring problem is, in the general case, universally insurmountable (assuming 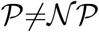) and (2) RVnRS does not endow the organism with any heuristic-based approximation shortcuts that offer better than brute-force exploration of the search space. A sparse MIN where a gene has at most one interacting partner produces the easiest possible instances of the network-rewiring problem: regardless of the nature of the current evolutionary pressure, any gene can either be beneficial or damaging depending on whether the interaction it is engaged in is beneficial or damaging to the network as a whole. There is therefore no ambiguity as to which genes to conserve and which to delete/mutate in such a fully sparse network. However, such networks would necessarily contain more genes, since functions that could have otherwise been accomplished by a single hub gene (e.g. a phosphatase targeting multiple proteins) must now be handled by a large number of specialty genes. This clearly leads to an explosion of genome size. On the other extreme, a dense network where functions are concentrated in as few multi-tasking hub genes as possible would lead to an exponential search space. Particularly, the number of iterations of random-variation non-random selection needed before the network has been re-wired away from a deleterious and into an advantageous state would be exponential in the number of unambiguous genes given some evolutionary pressure. That all genes in such a network are engaged in more than one interaction exponentially decreases their likelihood of being unambiguously advantageous or disadvantageous for the organism under a given evolutionary pressure scenario. An exponential number of iterations of random-variation and non-random selections are needed before the right set of genes have been conserved, deleted or mutated such that the total number of damaging interactions network-wide is under a tolerable threshold.

mLmH topology is the middle ground between the two aforementioned extremes (the totally sparse but large, and the highly dense but unadaptable networks): essential functions are concentrated [34] in ‘hub’ genes that are unlikely to be damaging in and of themselves [36] while continuous (and cheap) experimentation (e.g. fine-tuning micro-RNA regulation [34]) is conducted at the periphery of the network [37] with loosely connected leaf genes.

### applications and extensions

An immediate application of the model is to complement statistical tests used to infer the quality and coverage of large-scale interactome-mapping wet experiments [1] or *in-silico* network inference [38], by testing whether the resulting networks over- or under-represents real interactions relative to the prediction. There are aspects of the model that should be extended. In this work we treated all interactions as equal, but in reality some interactions are more potent than others. Future work could extend the model by considering the potency of each interaction, a not so trivial task since no large-scale data exist yet to facilitate its inference. The model is also static, in that assigning benefit/damage to a gene is based on immediate neighbours only. The implication is that all genes are equal, but in reality a central gene (many shortest paths pass through it) has much more effect network-wide than a gene residing at the periphery of the network. Trivially, implementing a dynamic variant (where the cascading effect of a gene’s beneficial (damaging) effects are accounted for) does not change the complexity class of NEP, although its simulations will be more computationally demanding.

## Abbreviations

NPC: 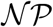-complete
MIN: Molecular interaction network
mLmH: majority-Leaves minority-Hubs network topology
OA: Oracle Advice
RVnRS: Random variation non-random selection
NEP: Network Evolution Problem
KOP: Knapsack Optimization Problem
PPI: Protein-protein interaction
NL: No-Leaf network
NH: No-Hub network
*amb*: ambiguous
EIS: Effective instance size
PSICQUIC: Proteomics Standard Initiative Common QUery InterfaCe
*e*2*n*: edge:node ratio of a network

## Supplementary materials

Supplementary Text

Figs. S1 to S17

Tables S1 - S4

## Competing interests

the Authors declare no competing financial or non-financial interests.

## Author contribution

AAA, KI, SV and JW designed research; AAA performed research, implemented computer simulations, analyzed data, and wrote the paper; CH implemented simulated evolution experiments; JW, KI, SV, CH revised and corrected the paper.

## Data availability

The data/reanalysis that support the findings of this study are publicly available online at: http://cs.mcgill.ca/malsha17/permlink/NETWORKS/.

## Code availability

Model codes are available at https://github.com/aliatiia/EvolByCompSel.

## Acknowledgements

Computations were made on the supercomputing cluster Guillimin from McGill University, managed by Calcul Québec and Compute Canada. The operation of these supercomputers is funded by CFI, MESI, and FRQ-NT.

## Footnotes

1 mLmH-possessing networks are typically referred to as ‘scale-free’ (SF). Because of the association between the SF term and the controversial proposition [5, 6, 7, 8] that biological networks follow a power-law distribution, we use the loose term mLmH to emphasize the fact that our model does not assume, require nor advocate for or against the idea that the degree distribution of MINs follows a power-law in a strict technical sense.

2 These figures are the average over the 25 networks featured in this article. To our knowledge, these networks represent all large-scale experimentally validated networks of direct physical interactions available in the literature.

## Notes

#### Summary of Updates

slight amendment of summary/abstract.

