## Supplementary information for "Computational Intractability Law Molds the Topology of Biological Networks"

### Supplementary Material

Ali Atiia,[1,2] Corbin Hopper[1], Katsumi Inoue[3], Silvia Vidal[2], Jérôme Waldispühl[1*]
[1]School of Computer Science, McGill University, Montreal, Canada
[2]Research Centre on Complex Traits, McGill University, Montreal, Canada
[3]National Institute of Informatics, Tokyo, Japan

#### 1 Abbreviations

| | |
|---|---|
| **NPC** | $\mathcal{NP}$-complete |
| **MIN** | Molecular interaction network |
| **mLmH** | majority-Leaves minority-Hubs network topology |
| **OA** | Oracle Advice |
| **RVnRS** | Random variation non-random selection |
| **NEP** | Network Evolution Problem |
| **KOP** | Knapsack Optimization Problem |
| **PPI** | Protein-protein interaction |
| **NL** | No-Leaf network |
| **NH** | No-Hub network |
| *amb* | ambiguous |
| **EIS** | Effective instance size |
| **EGB** | Effective gained benefits |
| **PSICQUIC** | Proteomics Standard Initiative Common QUery InterfaCe |
| **MIQL** | Molecular Interaction Query Language |
| $n2e$ | node:edge ratio of a network |
| $e2n$ | edge:node ratio of a network |

### 2 MINs with experimental evidence and their synthetic analogs:

#### 2.1 protein-protein interaction networks

Table 1 shows the details and references of protein-protein interaction (PPI) networks (raw data and source code available in [1]). PPI networks represent a "universe of possibilities", where combinatorial experiments test the affinity of each protein against all others in (typically, in large-scale experiments) exogenous settings. Widely used experimental methods include yeast two-hybrid (Y2H) and affinity purification followed by Mass spectrometry (AP-MS). Examining the literature references in Table 1 in chronological order of publication dates (ranging from 2008-2016), one observes a rapid increase in the scale and resolution of high-throughput methods with works by Rolland et *al.* [2] and Yang et *al.* [3] representing the cutting edge in terms of coverage and resolution respectively. In [3], it was shown that different isoforms of the same protein can exhibit quite different interaction profiles. Therefore the degree of a gene (particularly hub genes) may in fact be inflated in networks where isoforms are not distinguished: that gene should ideally be broken down to separate nodes corresponding to each isoform. Typically, further validation of the resulting networks is conducted on a subset of interactions by testing their affinity in endogenous setting (which in turn is used to calculate some measure of true/false positives/negatives or some combination of such ratios) or comparing the resulting interactions to (small) gold standard data sets. It is important to note that PPI networks are generally undirected, since the experimental methods only establish the existence of an interaction but reveal nothing about the type (whether promotional or inhibitory) or directionality (which of the two proteins affects the other) of an interaction. The Fly network is the one exception, as both the direction and type of its interactions have been assessed using a simple prediction algorithm which achieved "90% precision and 41% recall (2.8% false positive rate and 59% false negative rate)" [4]. Figure SI 1 shows the degree distribution of PPI networks and their corresponding synthetic analogs which were generated using the same method discussed in Section "Simulation of evolutionary pressure" in the main text.

| PPI Network | no. nodes | no. edges | e2n ratio | directed? | signed? |
|---|---|---|---|---|---|
| Plant [5] | 2661 | 5664 | 2.13 | no | no |
| Bacteria [6] | 1267 | 2233 | 1.76 | no | no |
| Yeast [7] | 2018 | 2930 | 1.45 | no | no |
| Worm [8] | 2528 | 3864 | 1.53 | no | no |
| Fly [4] | 3352 | 6094 | 1.82 | yes | yes |
| Human [2] | 4303 | 13944 | 3.24 | no | no |
| HumanIso [3] | 629 | 996 | 1.58 | no | no |

Table 1: Summary of protein-protein interaction (PPI) networks. The direction and sign of an interaction were assigned at random (coin flip) in undirected and/or unsigned networks. References, data and source code publicly available in [1].

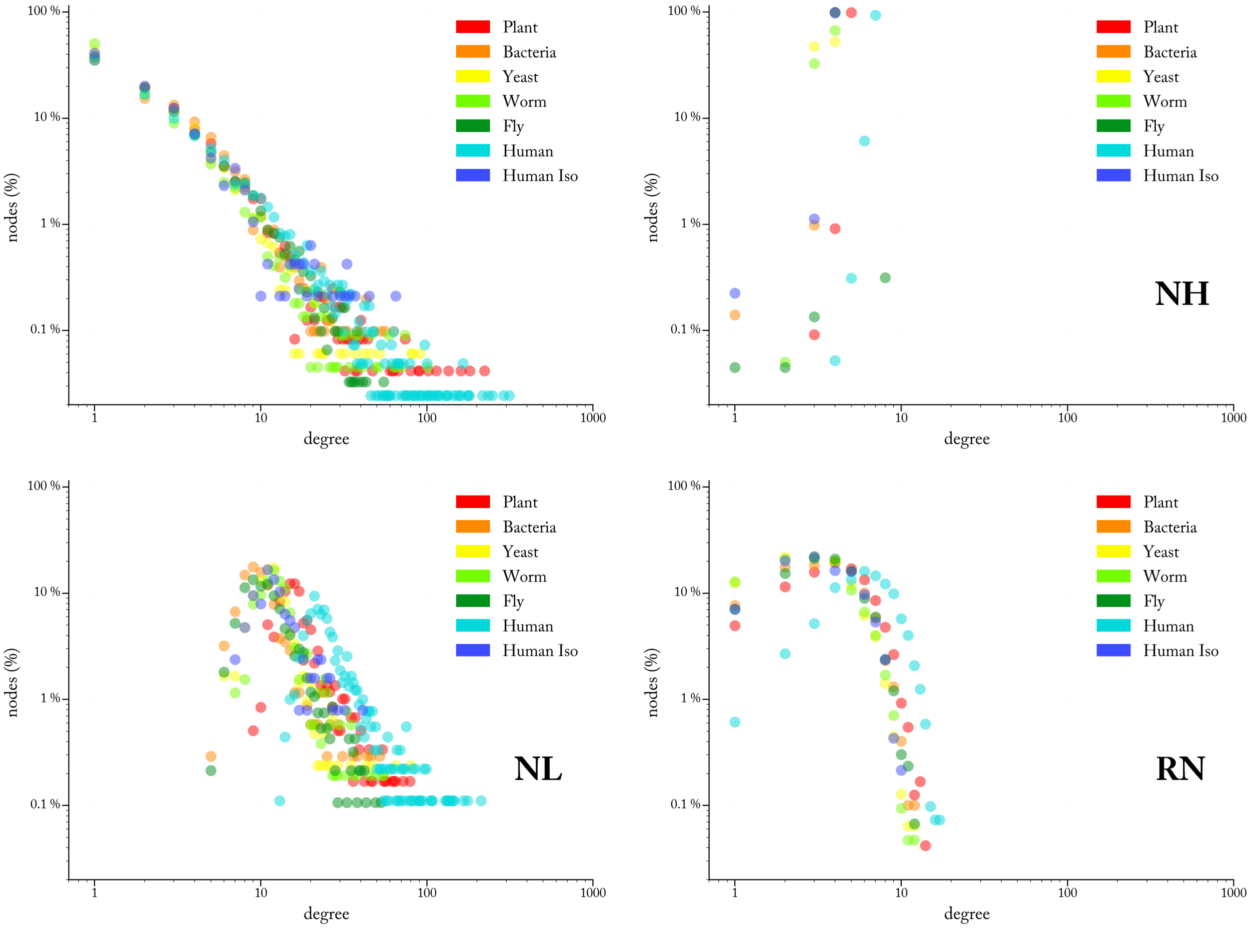


Figure 1: Degree distribution of PPI networks and their corresponding synthetic analogs: no-hubs (NH), no-leaves (NL) and random (RN).

#### 2.2 Regulatory networks

Regulatory networks (details and references in Table 2, raw data and source code available in [1]) are all directed, with some being partially signed (RegulonDB and TRRUST). The nodes in regulatory networks can be transcription factors, genes (which can refer to the protein or mRNA), or small RNAs. All networks originally contain exclusively experimentally-validated interactions, with the exception of Liu and RegulonDB which contain computationally (*in-silico*) inferred interactions which were excluded. In the case of RegulonDB, only interactions with 'strong' or 'confirmed' experimental evidence are included, and since none of the interactions involving small RNAs had such evidence, they were eliminated. The remaining interactions were therefore exclusively between transcription factors. In miRTarBase networks, only interactions with strong experimental evidence (elucidated through reporter assays or western blot experiments) are included. Furthermore, interactions where the species of source and target genes are different were excluded (presumably, these original from transgenic studies).

The ENCODE proximal network is an overall consolidated network of transcriptional interactions in humans, with some interactions being obtained by further consolidation with PPI network (detailed in supplementary materials of [9]). The other two ENCODE networks on the other hand are generated from specific human cell lines (GM and K562). The TRRUST network is unique in that it was obtained by data mining ∼20 million literature abstracts from Medline (2014), out of which ∼23K sentences were nominated to contain potential descriptions of regulatory interactions [10]. These sentences underwent successive rounds of manual inspections. TRRUST network also includes information about the nature of interactions and the number of studies supporting it. For interactions deemed promotional by some studies and inhibitory by others, we picked the sign randomly by flipping a crooked coin proportional to the number of studies that support one type or another (for example, if 3 studies report an interaction as 'promotional' and 1 reports it as 'inhibitory', we would consider the interaction to be 'promotional' with 75% likelihood). TRRUST authors aimed to create a high-quality network that can serve as a gold-standard to other large-scale studies aiming to map transcriptome interactions in humans. The same crooked coin strategy was used in RegulonDB network. Figure SI 2 shows the degree distributions of regulatory networks and their corresponding synthetic analogs. Despite the diverse methods that were behind the mapping of these networks (in contrast to PPIs, where Y2H method is dominant), the mLmH property still holds with lower-degree nodes in particularly being of almost the same frequency in the majority of networks.

| Regulatory Network | no. nodes | no. edges | e2n ratio | directed? | signed? |
|---|---|---|---|---|---|
| Bacteria RegulonDB [11] | 898 | 1481 | 1.649 | yes | no |
| ENCODE Proximal [9] | 9057 | 26070 | 2.878 | yes | no |
| ENCODE K562 [9] | 3947 | 9595 | 2.431 | yes | no |
| ENCODE GM [9] | 3989 | 6971 | 1.748 | yes | no |
| Human Liu [12] | 3502 | 9606 | 2.743 | yes | no |
| Human TRRUST [10] | 2718 | 8015 | 2.949 | yes | yes |
| Human miRTarBase [13] | 2583 | 5450 | 2.11 | yes | no |
| Mouse Liu [12] | 1436 | 3673 | 2.558 | yes | no |
| Mouse miRTarBase [13] | 741 | 1019 | 1.375 | yes | no |

Table 2: Summary of regulatory networks. The direction and sign of an interaction were assigned at random (coin flip) in undirected and/or unsigned networks. References, data and source code publicly available in [1].

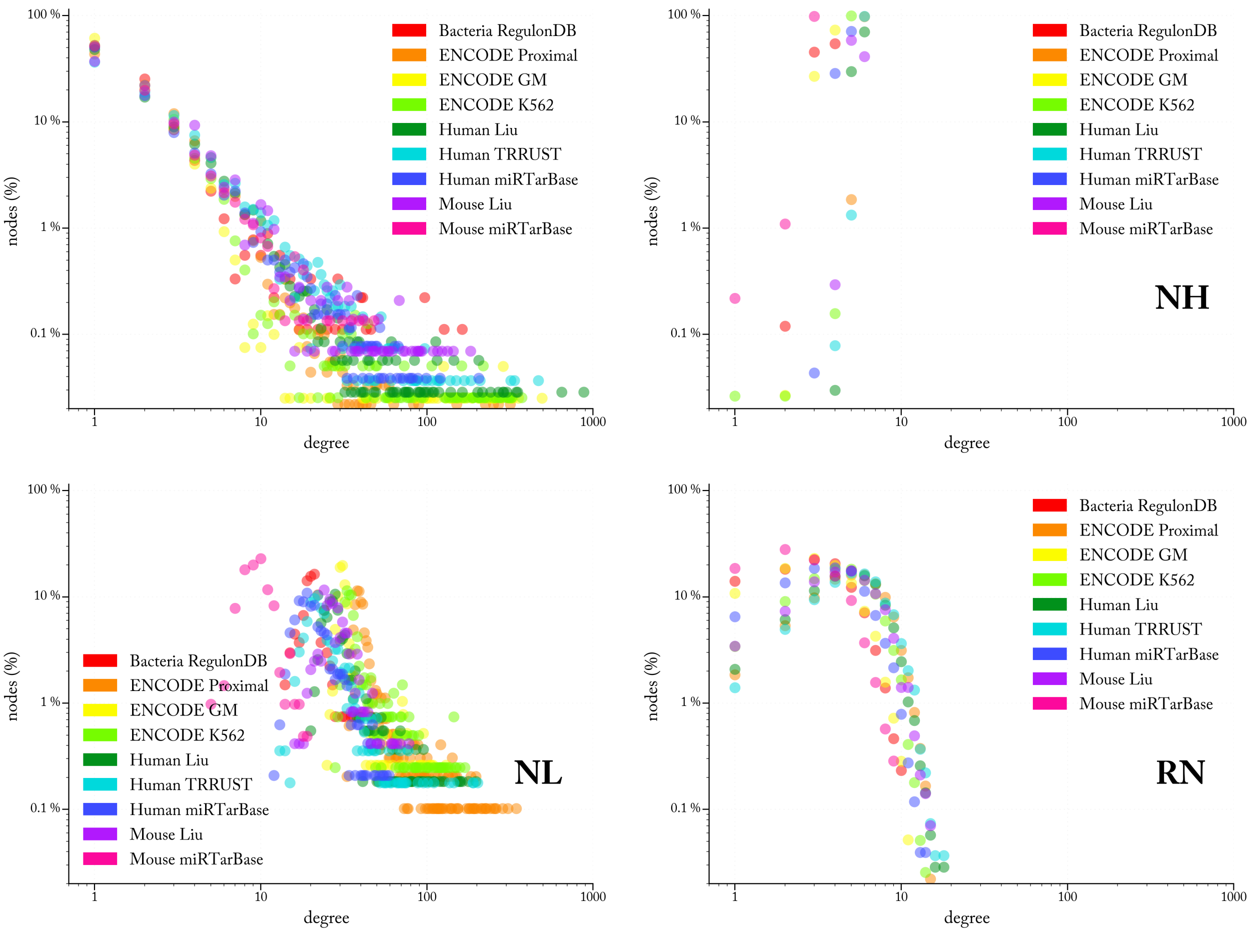


Figure 2: Degree distribution of regulatory networks and their corresponding synthetic analogs: no-hubs (NH), no-leaves (NL) and random (RN).

#### 2.3 Database-sourced networks

Table 3 shows details and source references of networks obtained from the BioGrid database or from multiple databases queried simultaneously through the PSICQUIC web service (raw data and source code available in [1]). All obtained interactions are undirected and unsigned. Interactions in BioGrid networks represent physical interactions which have been validated by at least two studies, except for human and yeast networks in which interactions have been validated by at least 4 and 3 studies, respectively (because of the large number of interactions for these two species, it was still possible to obtain large networks even under this stringent selection criteria). Multiple databases (excluding BioGrid) were searched programmatically with a Molecular Interaction Query Language (MIQL) query through the PSICQUIC web service interface (source code publicly available in [1]). The query specifies interactions where both interactors (1) are from the same species, (2) they interact physically, and (3) the interaction has been experimentally detected. It should be noted that some PSICQUIC interactions did distinguish whether an interactor is an isoform of a well-known gene. Figure SI 3 shows the degree distribution of the resulting networks and their corresponding synthetic analogs. The Plant-PSICQUIC network is anomalous in its degree distribution, indicating sporadic coverage of its reported interactions. Other networks of even smaller size still exhibit the mLmH property, which can be a sign that the underlying studies behind them were less sporadic in their coverage (i.e. focusing on specific functional units).

| DB-sourced Network | no. nodes | no. edges | e2n ratio | directed? | signed? |
|---|---|---|---|---|---|
| Plant-BioGrid [14] | 1565 | 2745 | no | no | |
| Plant-PSICQUIC [15] | 230 | 789 | 3.43 | no | no |
| Yeast-BioGrid [14] | 2418 | 7668 | 3.171 | no | no |
| Yeast-PSICQUIC [15] | 767 | 1386 | 1.807 | no | no |
| Worm-BioGrid [14] | 55 | 64 | 1.164 | no | no |
| Fly-BioGrid [14] | 188 | 279 | 1.484 | no | no |
| Mouse-BioGrid [14] | 1031 | 1497 | 1.452 | no | no |
| Human-BioGrid [14] | 3436 | 8254 | 2.402 | no | no |
| Human-PSICQUIC [15] | 3470 | 6188 | 1.783 | no | no |

Table 3: Summary of real database-sourced networks. The direction and sign of an interaction were assigned at random (coin flip) in undirected and/or unsigned networks. References, data and source code publicly available in [1].

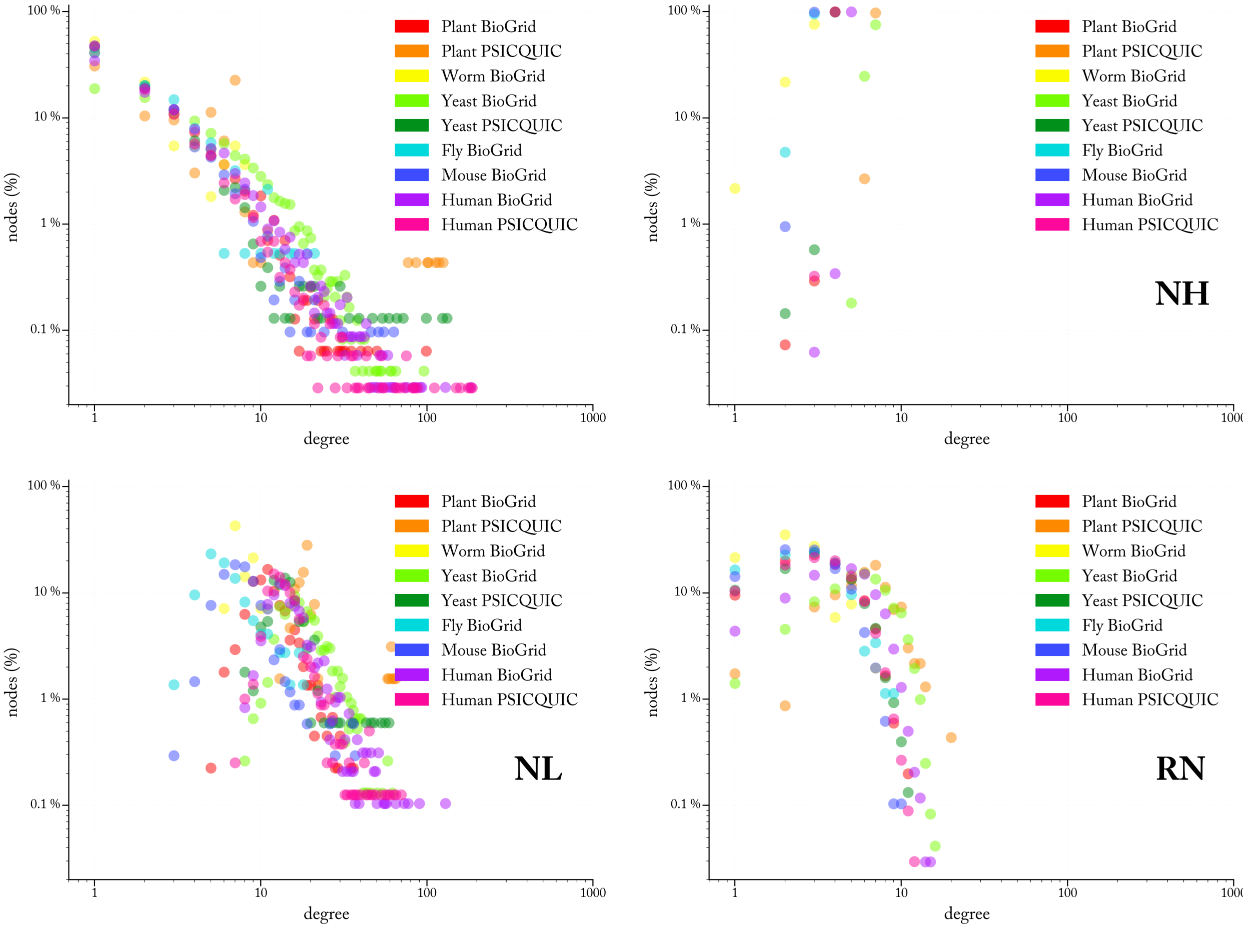


Figure 3: Degree distribution of DB-sourced networks and their corresponding synthetic analogs: no-hubs (NH), no-leaves (NL) and random (RN).

### 3 Formal Definition of the Network Evolution Problem (NEP)

Given:

$$\boldsymbol{G} = (g_1, g_2, \ldots, g_n),\ \boldsymbol{A} = (a_1, a_2, \ldots, a_n),\ a_j \in \{+1, 0, -1\},\ \ \boldsymbol{t} \in \mathbb{R},\ \text{ and}$$
$$\boldsymbol{M} = \left[m_{jk}\right] \quad \text{where} \quad m_{jk} \in \mathbb{R}, \quad \forall j,k,\ 1 \leq j,k \leq n$$

Let:

$$\boldsymbol{B} = (b_1, b_2, \ldots, b_n),\ \text{where}\ \ b_j = \sum_{k=1}^{n} m_{jk} \oplus a_k \ \ + \ \sum_{k=1}^{n} m_{kj} \oplus a_j \ \ \text{and}$$
$$m_{xy} \oplus a_y = \begin{cases} |m_{xy}| & \text{if} \quad m_{xy} \times a_y > 0 \\ 0 & \text{otherwise} \end{cases}$$

$$\boldsymbol{D} = (d_1, d_2, \ldots, d_n),\ \text{where}\ \ d_j = \sum_{k=1}^{n} m_{jk} \ominus a_k \ \ + \ \sum_{k=1}^{n} m_{kj} \ominus a_j \ \ \text{and}$$
$$m_{xy} \ominus a_y = \begin{cases} |m_{xy}| & \text{if} \quad m_{xy} \times a_y < 0 \\ 0 & \text{otherwise} \end{cases}$$

Define:

$$f : \boldsymbol{G} \rightarrow \{0,1\} \text{ maximizing } \sum_{j=1}^{n} f(g_j) \times b_j \ \text{ s.t. } \left( \sum_{j=1}^{n} f(g_j) \times d_j \right) \leq \boldsymbol{t}$$

Table 4 provides a summary of each element of NEP and its corresponding semantic interpretation in biological context (see also the main text for more on the semantics of NEP in biological context).

| | |
|---|---|
| $\boldsymbol{G} = (g_1, g_2, \ldots, g_n)$ | A sequence of $\boldsymbol{G}$enes: any transcribable element on the genome |
| $\boldsymbol{A} = (a_1, a_2, \ldots, a_n)$ | A ternary string representing an Oracle $\boldsymbol{A}$dvice: $a_j = \begin{cases} +1 & \implies g_j \text{ is advantageous} \\ -1 & \implies g_j \text{ is disadvantageous} \\ 0 & \implies \text{no opinion on } g_j \end{cases}$ |
| $\boldsymbol{M} = \left[m_{jk}\right]$ | $n \times n$ $\boldsymbol{I}$nteraction $\boldsymbol{M}$atrix: $m_{jk} = \begin{cases} p > 0 & \implies g_j \text{ promotes } g_k \\ q < 0 & \implies g_j \text{ represses } g_k \\ 0 & \implies g_j \text{ and } g_k \text{ don't interact} \end{cases}$ |
| $\boldsymbol{t} \in \mathbb{R}$ | $\boldsymbol{t}$olerance threshold on damages |
| $\boldsymbol{B} = (b_1, b_2, \ldots, b_n)$ <br> $\boldsymbol{D} = (d_1, d_2, \ldots, d_n)$ | Each gene $g_j$ has a corresponding benefit value $\boldsymbol{b_j}$ and damage value $\boldsymbol{d_j}$ given an Oracle advice on $g_j$ and all or some of its interaction partners. |
| $m_{xy} \oplus a_y = \begin{cases} \lvert m_{xy} \rvert & \text{if } m_{xy} \times a_y > 0 \\ 0 & \text{otherwise} \end{cases}$ | If the effect of $g_x$ on $g_y$ is in **agreement** with what the Oracle says $g_y$ should be (i.e. $m_{xy}$ and $a_y$ have the same sign), then increment $b_x$ by $\lvert m_{xy} \rvert$ |
| $m_{xy} \ominus a_y = \begin{cases} \lvert m_{xy} \rvert & \text{if } m_{xy} \times a_y < 0 \\ 0 & \text{otherwise} \end{cases}$ | If the effect of $g_x$ on $g_y$ is in **disagreement** with what the Oracle says $g_y$ should be (i.e. $m_{xy}$ and $a_y$ have different signs), then increment $d_x$ by $\lvert m_{xy} \rvert$ |
| $f : \boldsymbol{G} \rightarrow \{0, 1\}$ <br> maximizing: <br> $\sum_{j=1}^{n} f(g_j) \times b_j$ <br> subject to: <br> $\left( \sum_{j=1}^{n} f(g_j) \times d_j \right) \leq \boldsymbol{t}$ | The idealistic pursuit of enforcing an Oracle advice (OA) is complicated by the reality of network connectivity: <br> OA can be imposed by deleting every gene $\boldsymbol{g_i}$ where $a_i = -1$ and conserving every gene $\boldsymbol{g_j}$ where $a_j = +1$. However: **deleting $\boldsymbol{g_i}$** can inadvertently contribute to a violation of the OA if $g_i$ happens to be a promoter (repressor) of some $g_k$ that should in fact be promoted (repressed); and **conserving $\boldsymbol{g_j}$** can inadvertently contribute to a violation of the OA if $g_j$ happens to be a promoter (repressor) of some $g_k$ that should in fact be repressed (promoted). What subset of genes should be conserved/deleted (define $f$) such that the OA is supported by as many interactions as possible (the *maximize .. subject to..* clauses)? |

Table 4: The syntax (left column) and semantics (right) of the network evolution problem (NEP)

### 4 NP-hardness of NEP

The $\mathcal{NP}$-hard knapsack optimization problem (KOP) [16] is defined as: Given a sequence of objects $\boldsymbol{O} = (o_1, o_2, \ldots, o_r)$, values $\boldsymbol{V} = (v_1, v_2, \ldots, v_r)$, weights $\boldsymbol{W} = (w_1, w_2, \ldots, w_r)$, and a knapsack capacity $\boldsymbol{c}$ where $v_i, w_i, \boldsymbol{c} \in \mathbb{N}$, define:

$$f : \boldsymbol{O} \to \{0,1\} \text{ maximizing } \sum_{j=1}^{r} f(o_j) \times v_j \text{ s.t. } \left( \sum_{j=1}^{r} f(o_j) \times w_j \right) \leq \boldsymbol{c}.$$

**Theorem**: *NEP is $\mathcal{NP}$-hard by reduction from KOP.*

#### 4.1 Proof sketch

For a given KOP instance with $r$ items, create a graph with $r+1$ nodes: $n_1, n_2 \ldots, n_{r+1}$. Assume an OA where $a_i = +1 \; \forall a_i \in A$ except for $a_{r+1} = 0$. For each $v_i \in \boldsymbol{V}$, draw a $v_i$-weighted edge from $n_i$ to itself. Sort objects in $\boldsymbol{O}$ ascendingly by their respective weights in $\boldsymbol{W}$, call this sorted list $\boldsymbol{O'}$. $\forall w_i \in \boldsymbol{W}$, draw a $-w_i$-weighted edge from $n_i$ to $n_j$ where $o_j$ is the successor of $o_i$ in $\boldsymbol{O'}$. Because $n_j$ is attracting damaging interactions due to incoming edges from $n_i$, update its weight to $w_j - w_i$. For the last node $n_r$, draw $-w_r$-weighted edge from node $n_{r+1}$ to $n_r$. Because $n_{r+1}$ has zero-value, it's ruled out *a priori* from the solution vector.

#### 4.2 Proof

I. Define $\boldsymbol{\gamma} : \{1,..,r\} \to \{1,..,r\}$ s.t. $\forall i, 1 \leq i < r : w_{\gamma(i)} \leq w_{\gamma(i+1)}$
II. Let $\boldsymbol{G} = \boldsymbol{O} + \{o_{r+1}\}$, $\boldsymbol{t} = \boldsymbol{c}$, $\boldsymbol{A} = (a_1, \ldots, a_r, a_{r+1})$, where $a_{r+1} = 0$ and $\forall i \leq r, a_i = +1$
III. Let $\boldsymbol{M}$ be a $d \times d$ zero-matrix, $d = r+1$. Populate $M$ as follows:
1. Repeat for $i = 1$ to $i = r-1$:

$$\begin{aligned} j &\leftarrow \gamma(i) \quad &\text{and} \quad k &\leftarrow \gamma(i+1) \\ m_{jj} &\leftarrow v_j \quad &\text{and} \quad m_{jk} &\leftarrow -w_j \\ w_k &\leftarrow w_k - w_j \end{aligned}$$

2. $j \leftarrow \gamma(r)$, $m_{jj} \leftarrow v_j$, $m_{dj} \leftarrow -w_j$

IV. Calculate $\boldsymbol{B}$, $\boldsymbol{D}$ and define $\boldsymbol{f} : \boldsymbol{G} \to \{0,1\}$ (Section 3).
V. Return $(f(o_1), \ldots, f(o_r))$ as KOP's solution vector ■

Proof notation follows that in KOP (above) and NEP (Section 3) definitions.

#### 4.3 Reverse-Reducing NEP To KOP

While the KOP-to-NEP reduction proves the later to belong to the same complexity class as the former, NEP-to-KOP reduction allows the use of an existing well-known pseudo-polynomial dynamic-programming algorithm [17] to solve instances of the former. NEP can be reverse-reduced to KOP by setting $O = G, V = B, W = D$, and $c = t$.

### 5 Oracle advice on interactions

An Oracle advice (OA) over interactions (edges) rather than genes (nodes), can be represented by a matrix $A = \left[a_{jk}\right]$ where $a_{jk} \in \{+1, -1\}$ if $m_{jk} \neq 0$ and $a_{jk} = 0$ otherwise (recall $m_{jk}$ is the entry at row $j$ and column $k$ of the interaction matrix $M$, see NEP definition in Section SI 3). While $m_{jk} \neq 0$ describes what the effect of $g_j$ on $g_k$ actually is, $a_{jk}$ describes what that effect should *ideally* be. A beneficial (damaging) interaction is one where $m_{jk} \times a_{jk} = 1$ ($m_{jk} \times a_{jk} = -1$). In other words, an interaction is beneficial (damaging) if it is in agreement (disagreement) with what the Oracle says that interaction should ideally be. For example, assume $g_j$ inhibits $g_k$, i.e. $m_{jk} = -1$, but the OA is $a_{jk} = +1$, then $m_{jk} \times a_{jk} = -1$ implies the real effect disagrees with the ideal and the interaction is deemed damaging. The benefit (damage) score of each gene $g_j$, given a matrix OA, is the sum of beneficial (damaging) interactions that $g_j$ is *projecting* onto (out-edges) or *attracting* from (in-edges) other genes in a similar manner as those calculated under a string OA (Section SI 3).

NEP remains $\mathcal{NP}$-hard under a matrix OA. To prove this, we modify the proof in Section 4 as follows:

I. Define $\boldsymbol{\gamma} : \{1, .., r\} \rightarrow \{1, .., r\}$ s.t. $\forall i, 1 \leq i < r : w_{\gamma(i)} \leq w_{\gamma(i+1)}$
II. Let $\boldsymbol{G} = \boldsymbol{O} + \{o_{r+1}\}$, $\boldsymbol{t} = \boldsymbol{c}$.
III. Let $\boldsymbol{M}$ be a $d \times d$ zero-matrix, $d = r + 1$. Populate $M$ as follows:
1. Repeat for $i = 1$ to $i = r - 1$:

$$\begin{aligned} j &\leftarrow \gamma(i) \quad \text{and} \quad k \leftarrow \gamma(i+1) \\ m_{jj} &\leftarrow v_j \quad \text{and} \quad m_{jk} \leftarrow -w_j \\ w_k &\leftarrow w_k - w_j \end{aligned}$$

2. $j \leftarrow \gamma(r)$, $m_{jj} \leftarrow v_j$, $m_{dj} \leftarrow -w_j$

IV. Let $A$ be a $d \times d$ matrix where:

$$a_{jk} = \begin{cases} +1 & \text{if} \quad m_{jk} \neq 0 \\ 0 & \text{otherwise} \end{cases}$$

V. Calculate $\boldsymbol{B}$, $\boldsymbol{D}$ as follows:

$$b_j = \sum_{k=1}^{n} m_{jk} \oplus a_{jk} + \sum_{k=1}^{n} m_{kj} \oplus a_{kj} \quad \text{where:}$$

$$m_{xy} \oplus a_{xy} = \begin{cases} 1 & \text{if} \quad m_{xy} \times a_{xy} > 0 \\ 0 & \text{otherwise} \end{cases}$$

and similarly the damage score is:

$$d_j = \sum_{k=1}^{n} m_{jk} \ominus a_{jk} + \sum_{k=1}^{n} m_{kj} \ominus a_{kj} \quad \text{where:}$$

$$m_{xy} \ominus a_{xy} = \begin{cases} 1 & \text{if} \quad m_{xy} \times a_{xy} < 0 \\ 0 & \text{otherwise} \end{cases}$$

VI. Define $\boldsymbol{f} : \boldsymbol{G} \rightarrow \{0, 1\}$ (Section 3)
VII. Return $(f(o_1), \ldots, f(o_r))$ as KOP's solution vector ■

### 6 Simulating Evolutionary Pressure

The simulation has the parameter tolerance $t$, expressed as percentages of total edges, indicating the total number damaging interactions to be tolerated (equivalently, the knapsack capacity $c$ in the corresponding KOP instance). For each network, the simulation is carried out under maximum pressure (non-zero OA on every gene) against each $t \in 0.1, 1, 5\%$. Given a tolerance value $t$, a knapsack instance is generated from a given NEP instance by reversing the reduction; that is: $O = G, V = B, W = D$ and $c = t$. The simulation records the total benefit and damage of objects (=genes, recall $O = G$) added to the knapsack by the solver [17] for each round against a randomly generated Oracle advice on each gene. The simulation is repeated for 1-5K iterations (sampling threshold, see Section 7). Figure SI 4 summarizes the algorithmic workflow of the simulation.

**right**: Simulations are carried at a certain pressure. Maximum pressure is when the Oracle has a non-zero advice on all nodes. Some simulations were carried at lower pressure levels where the Oracle is indifferent to 25, 50, 0r 75% of genes. For each tolerance $t$ value, 1-5K simulation rounds are carried out. In each round, a random OA is generated on all genes (nodes), followed by a calculation of benefit/damage value for each node against the current OA. The resulting NEP instance is reverse-reduced to a KOP instance ($O = G_i, V = B_i, W = D_i, c = t_i$) and fed to a knapsack solver. In each round, the sequences $G_i, B_i, D_i, t_i$, and $S_i$ are written to file, where $S_i$ is the solution vector $(s_1, \ldots, s_k), k = |G_i|$, and $s_i \in \{0, 1\}$. $s_i = 1$ ($s_i = 0$) implies "conserve" ("delete") or, in the context of the knapsack problem, "inside" ("outside") the knapsack.
**below**: average algorithm running time in milliseconds for each network. 'S' denotes an identical simulation on a second computer cluster different from the first run. For $t$=0.1%, the execution times are too negligible as a result of the dynamic programming algorithm [17] being upper-bounded by an exponent $= O(c)$ value. We therefore carried out the simulation at higher tolerance values $t \in \{5, 25, 50\}\%$. NL has significantly less nodes compared to other networks, and therefore shows the smallest execution times. PPI, RN and NH have ∼equal network sizes, but instances in PPI are solved faster compared to to its smaller instance sizes (a majority of genes being having either benefit (damage) as zero, and therefore such genes are not part of the optimization search as they should be conserved (deleted) regardless, see discussion on effective instance size (EIS) in the main text for details).

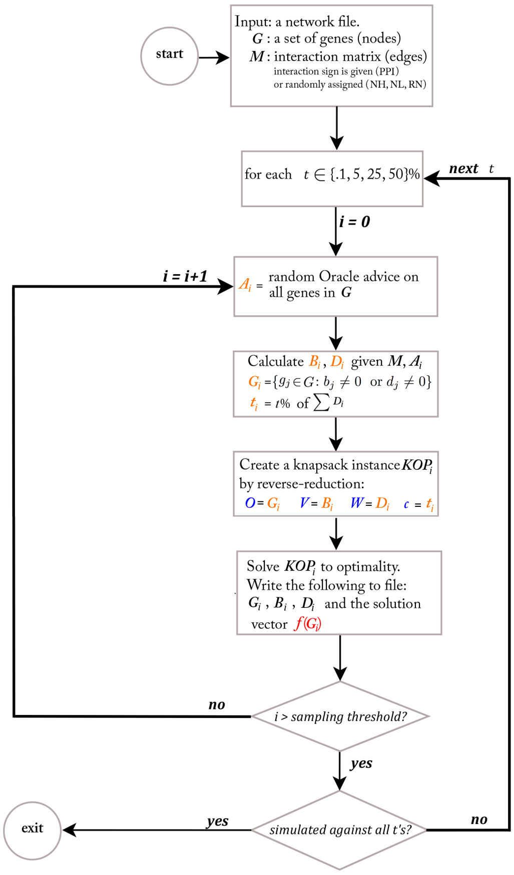


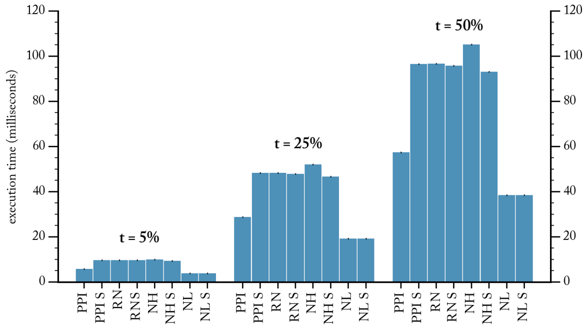


Figure 4: The algorithmic workflow of computer simulation and the average run time of the knapsack solver.

### 7 Effect of Sampling Threshold

Increasing the sampling threshold in the simulation (i.e. how many NEP instances to simulate) does not change the results, due to the effect of the Central Limit Theorem [18]. Figures SI 5 and 6 compare the results computed over 1,000 versus 5, 000 simulated instances (see the corresponding Figures 4 and 5 in the main text for detailed description).

#### 7.1 Benefit-Damage Correlation:

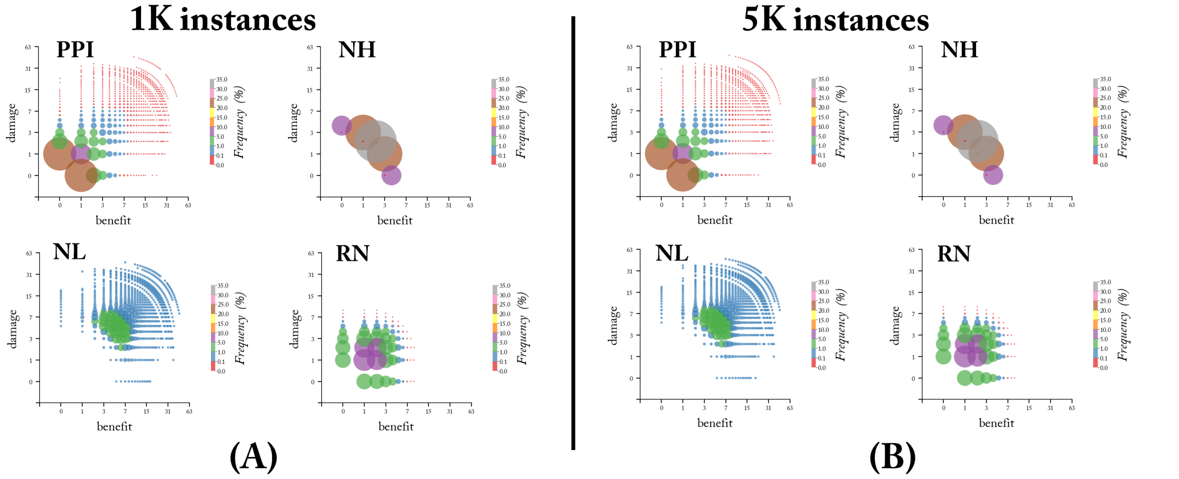


Figure 5: Increasing the sampling threshold from (A) 1,000 to (B) 5,000 NEP has virtually no effect on the resulting benefit-damage correlations.

#### 7.2 Effective Instance Size:

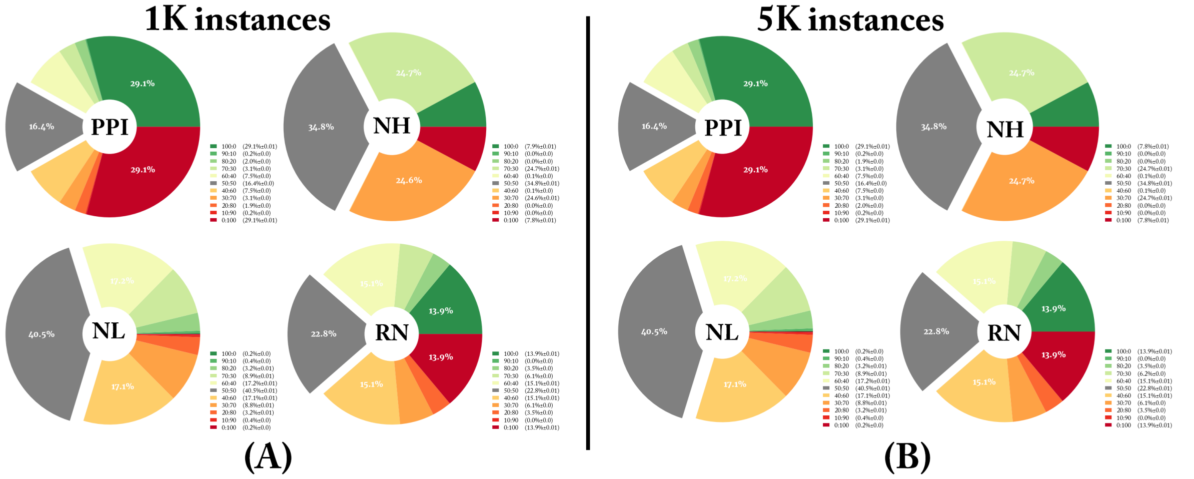


Figure 6: Increasing the sampling threshold from (A) 1,000 to (B) 5,000 NEP has minimal to no effect on effective instance size (EIS). Legend: numbers between parenthesis are average +/- standard deviation.

### 8 Benefit-damage correlation

Figures SI 7 and 8 show the benefit-damage correlation results for the regulatory and DB-sourced networks, respectively. For detailed description please see Figure 4 in the main text which shows the results for PPI networks.

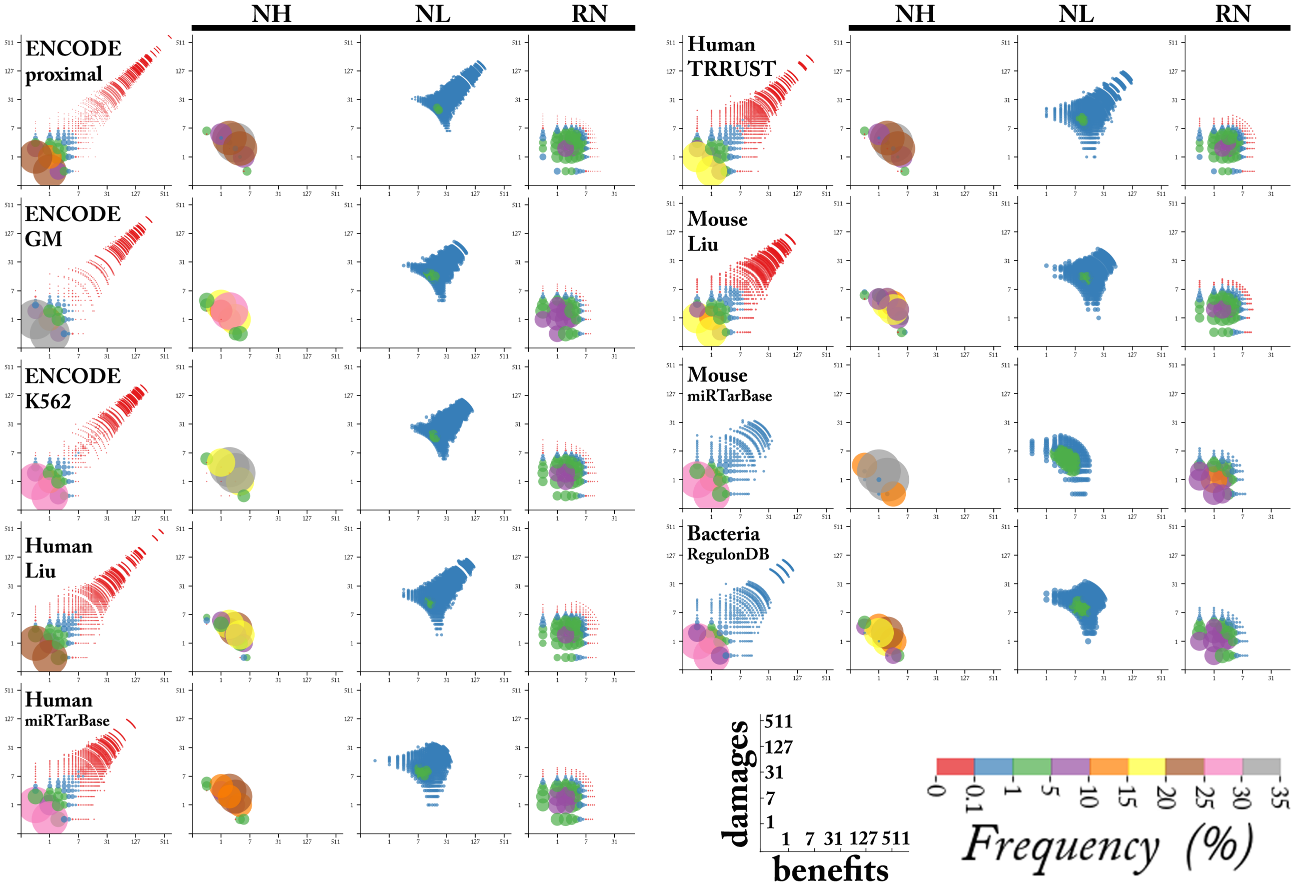


Figure 7: benefit-damage correlation in regulatory networks.

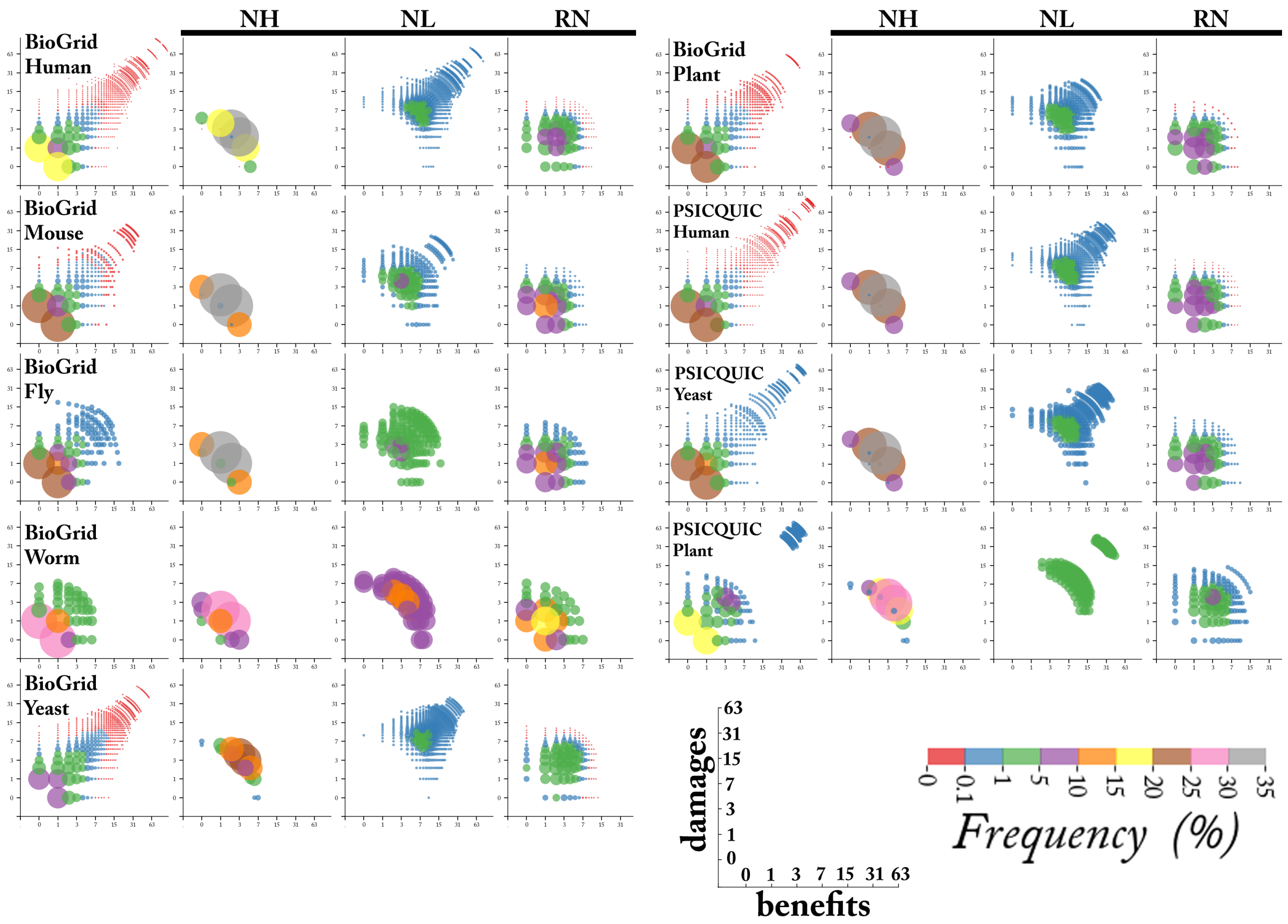


Figure 8: benefit-damage correlation in database-sourced networks.

### 9 Predicted vs. actual degree distributions

Figures SI 9, 10 and 11 show detailed plots of the actual versus predicted degree distribution of PPI, regulatory and DB-sourced networks, respectively, along with detailed bar plots of each $(\alpha, \beta)$ values used in the prediction formula and their respective proportionality to the node:edge ($n2e$) and edge:node ($e2n$) ratios in each networks. The $(\alpha, \beta)$ values were numerically determined by considering each $\alpha$ in the interval [0.01, 1] in increments of 0.01 against each $\beta$ in [0.1, 10] interval in increments of 0.1. Hub prediction may visually appear to be less precise but that is only due to the log scale in the y-axis. High discrepancies between $(\alpha, \beta)$ and $(n2e, e2n)$ values can be used to infer the quality of coverage and resolution of a network, and the extend to which it represents a representative sample the overall true and complete network. For example, $e2n >> \beta$ for the Yeast BioGrid network (Figure SI 11, right bar plot). Examining the degree distribution of this network (Figure SI 3), the frequency of degree-1 nodes is significantly low (~19%) compared to all other networks (DB-sourced, regulatory or PPI networks, where degree-1 frequency is 44±10%). The Worm BioGrid network on the other hand, has $\beta >> e2n$, which can be explained by the under representation of hub nodes in its network (it has no genes of degree $\geq$ 9, while on average 8±5% of genes in other networks have degree $\geq$9).

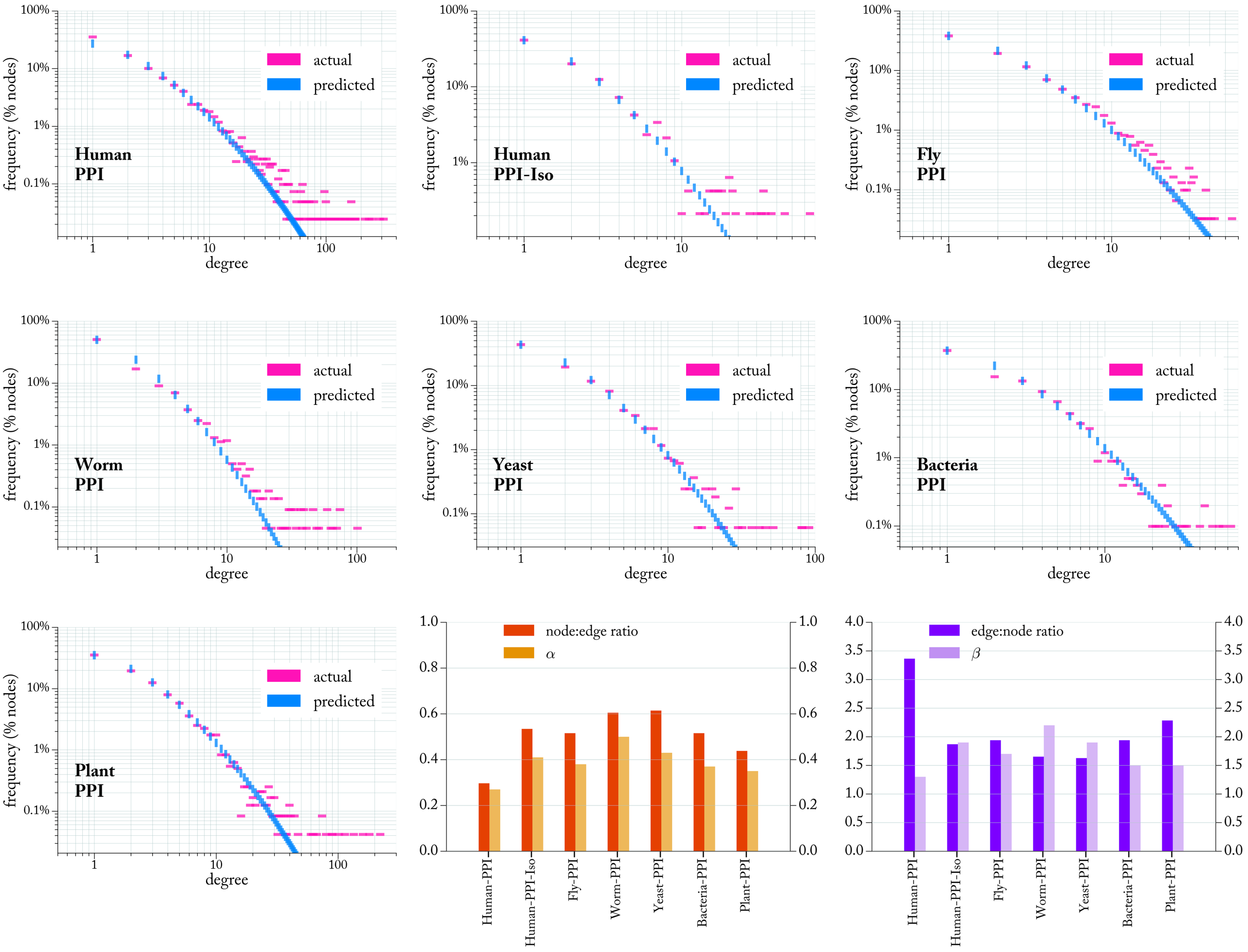


Figure 9: Actual and predicted degree distribution of PPI networks. The bar plots (bottom) show the $\alpha$ and $\beta$ values in the predicted networks versus the node:edge ($n2e$) and edge:node ($e2n$) ratios of the real networks.

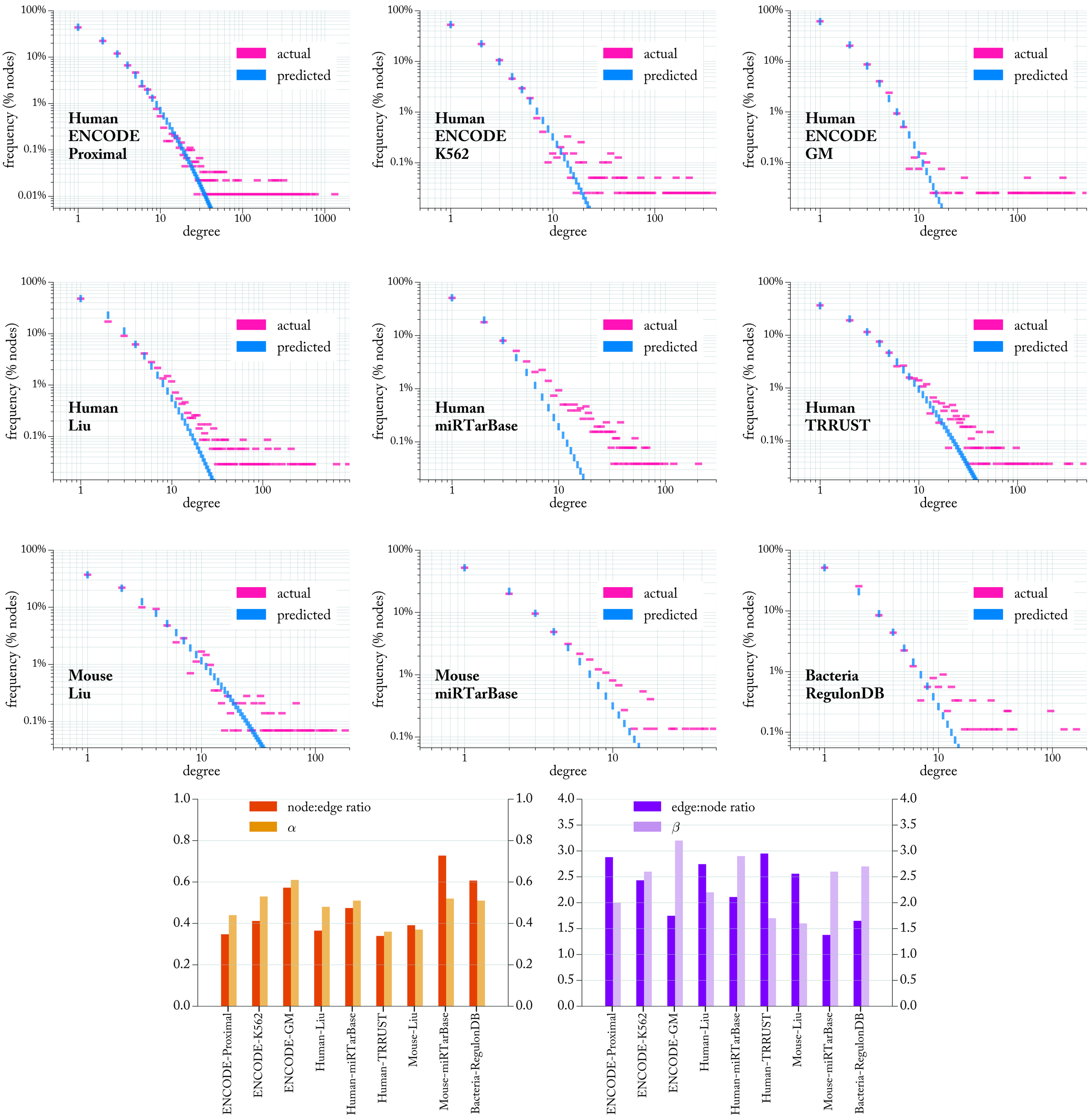


Figure 10: Actual and predicted degree distribution of regulatory networks. The bar plots (bottom) show the $\alpha$ and $\beta$ values in the predicted networks versus the node:edge ($n2e$) and edge:node ($e2n$) ratios of the real networks.

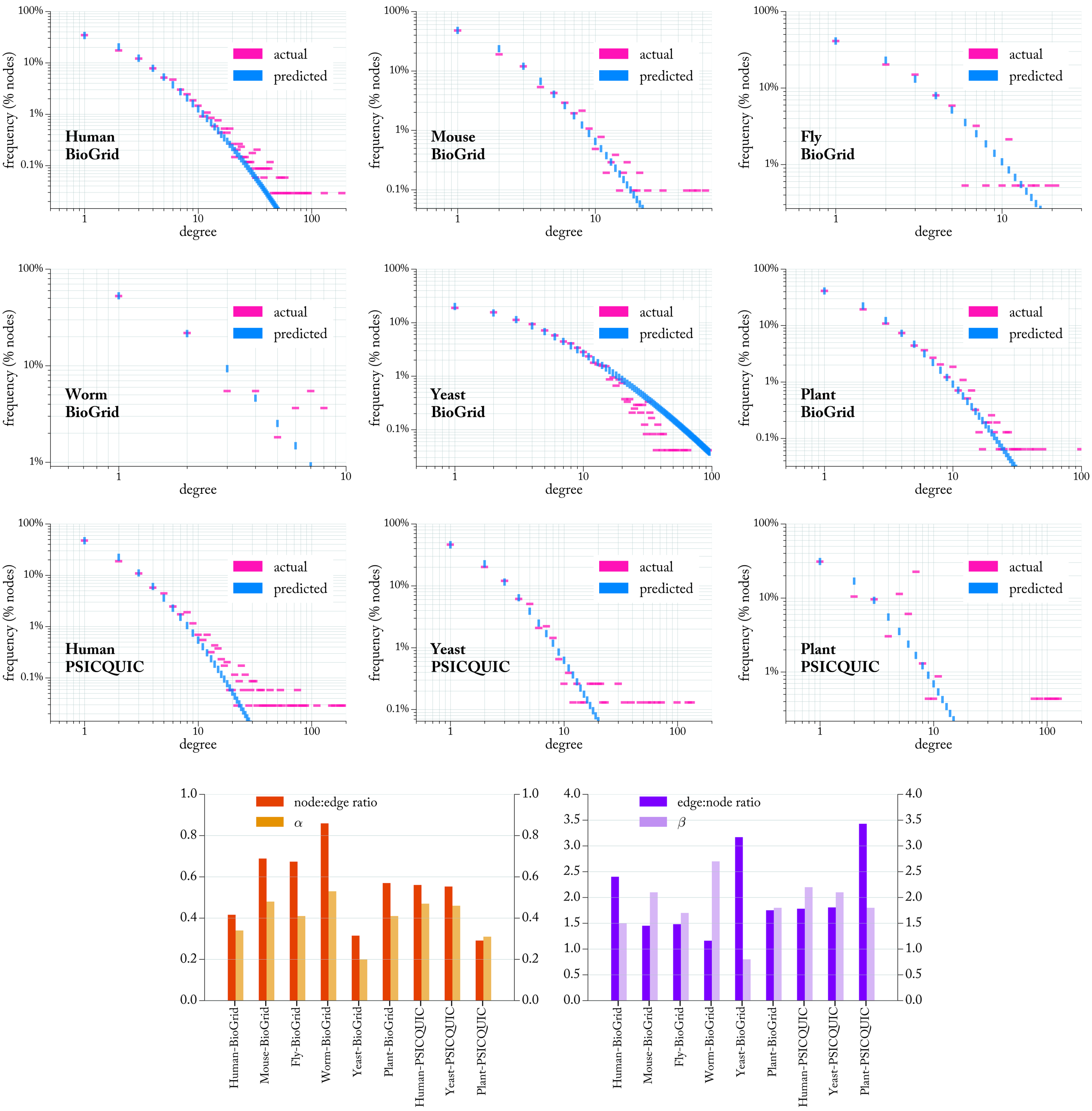


Figure 11: Actual and predicted degree distribution of database-sourced networks. The bar plots (bottom) show the $\alpha$ and $\beta$ values in the predicted networks versus the node:edge ($n2e$) and edge:node ($e2n$) ratios of the real networks.

### 10 Simulated evolution

Figure SI 12 shows the algorithmic workflow of the simulated evolution. In the fitness calculation step (denoted by a star in Figure SI 12), the fitness of a network vis-a-vis the current NEP instance is calculated based on 1) effective instance size (EIS) and 2) effective gained benefits (EGB) [19]. EIS is defined as the % of unambiguous nodes in the instance:

$$EIS = \frac{|\{n_i : b_i = 0 | d_i = 0\}|}{N} \tag{1}$$

where $N$ is the total number of nodes and $b_i$ and $d_i$ and the benefit and damage score of some node $n_i$. Let $(s_1, s_2, .., s_k)$ be the solution vector to an NEP instance where $s_i \in \{0, 1\}$ and $s_i = 1$ ($s_i = 0$) implies "conserve" ("delete"). The multiset $B = \{b_i : s_i = 1\}$ contains a list of all the benefits of nodes that are to be optimally conserved, and the effective gained benefits EGB=$sum(set(B))$. EGB is hence $B$ normalized by the number of nodes it takes to add a certain benefit value). For example, with $B1 = \{1, 1, 1, 5\}$ and $B2 = \{2, 6\}$, $sum(B1) = sum(B2) = 8$. But $EGB1 = sum(set(B1)) = sum(1, 5) = 6$ while $EGB2 = sum(set(B2)) = sum(2, 6) = 8$. Let $B_{tot}$ be the total benefit in a given NEP instance (the sum of gained benefits of conserved genes and lost benefits of deleted genes), the fitness of a given NEP instance $S$ is measured as:

$$F(S) = EIS^{\alpha} \times \frac{EGB}{B_{tot}} \tag{2}$$

where $\alpha \in\approx \mathbb{R}^+$. We applied $\alpha = 2$ in all simulation. With $\alpha$, the weight given to EIS vs EGB can be calibrated. This reflects the inherent opposition of EIS vs EGB, as EIS is best minimized with a large number of leaves while EGB is maximized with a large number of hubs. While EGB indicates how well a network accumulates as many beneficial interactions as possible with the smallest possible number of genes to conserve, normalizing it by the total benefits $B_{tot}$ penalizes networks that hemorrhage beneficial interactions that are lost to deleted genes in the optimal solution. The threshold $t$ of tolerated damaging interactions in the solution is imposed at 5% of the sum of all damages in all simulations.

Figures SI 13, 14 and 15 show extended results of the degree distribution of synthetically evolved networks grouped by network families (PPIs, regulatory and DB-sourced networks respectively). In each simulation, the network growth (add-edge and add-node mutations) are halted when the size of the synthetic network has become equal to the corresponding real network. Once network growth is disabled, the evolutionary algorithm is further run for a constant 2000 generations with re-assign edge mutation only.

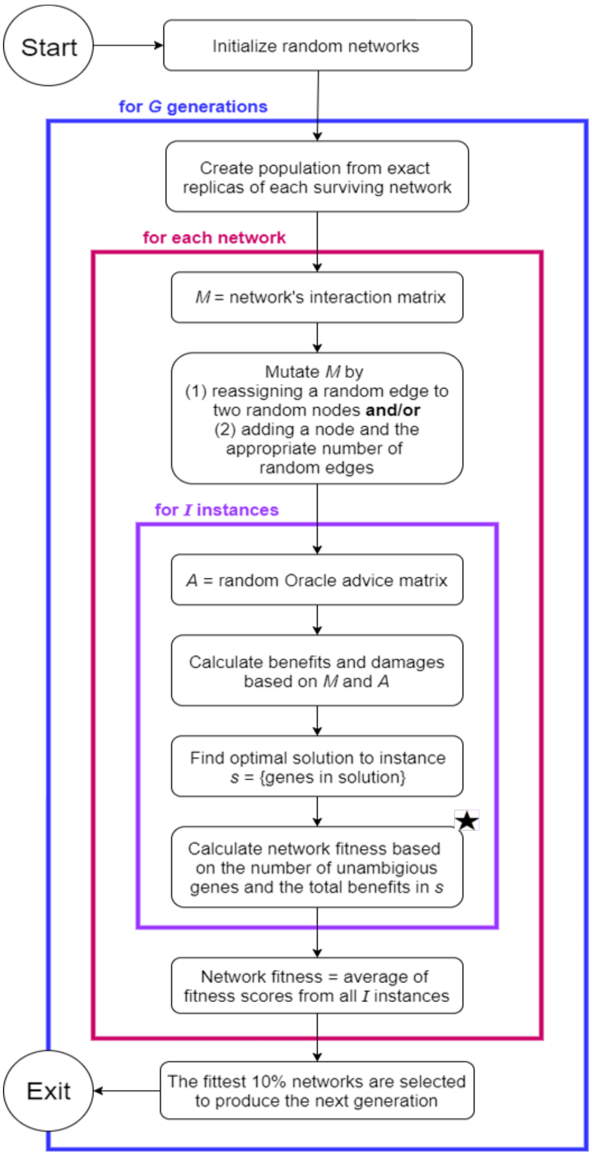


Figure 12: The algorithmic workflow of the evolutionary algorithm. Simulations begin with empty networks or seed networks that have randomly distributed edges. Each network is randomly mutated by reassigning one edge at each generation and, if growth is allowed, one node is also added along with as many randomly assigned edges as needed to maintain the desired edge:node ratio. An instance of the network evolution problem (NEP) is obtained by generating a random Oracle advice (OA) on all edges in the network. A network's fitness at each instance $S$ is calculated following the $F(S)$ formula (see text). The 10% of networks with the highest average fitness over all instances are selected to breed a population of networks for the subsequent generation. Adapted with modification with permission from [19].

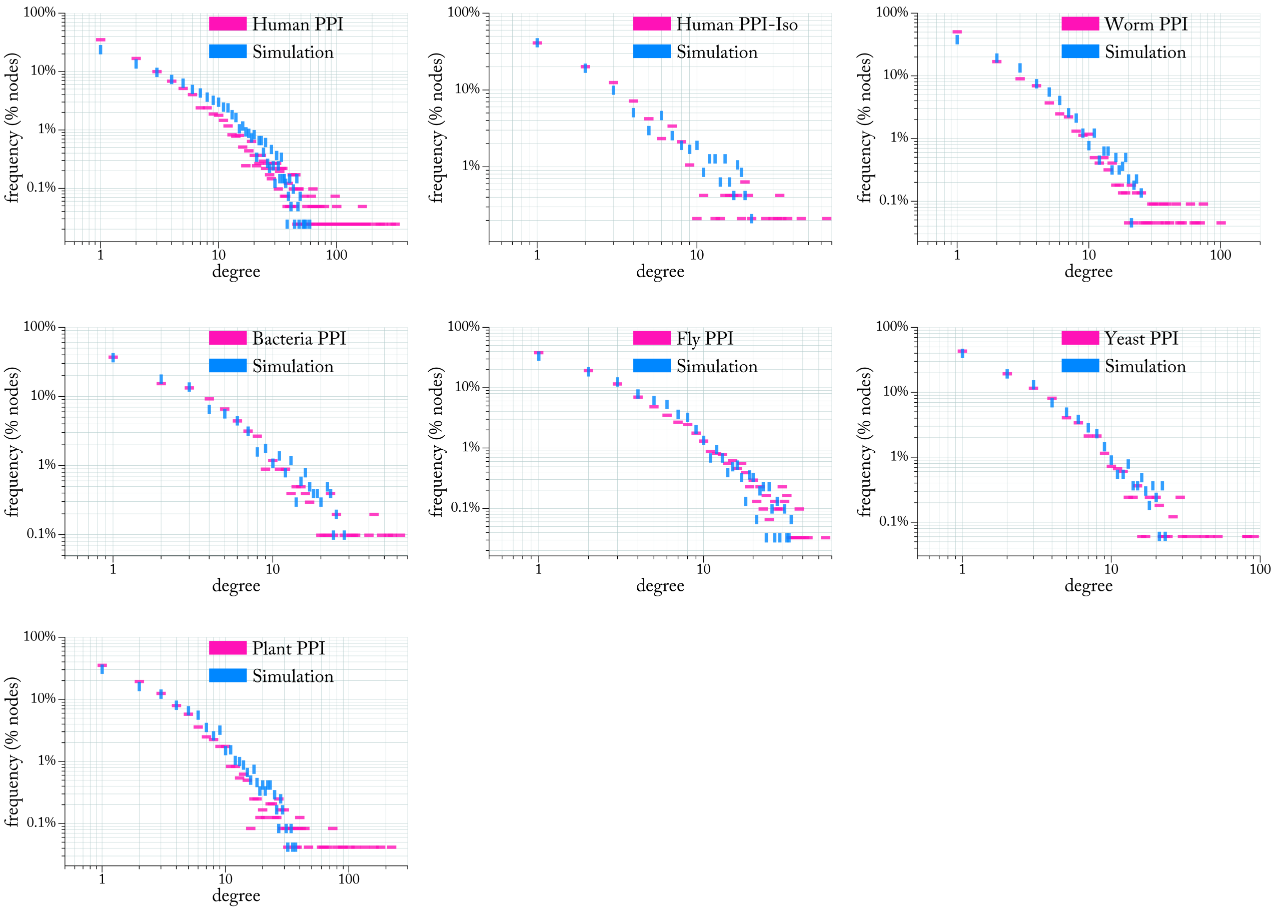


Figure 13: Evolving synthetic networks to the same size (number of nodes and edges) as protein-protein interaction networks.

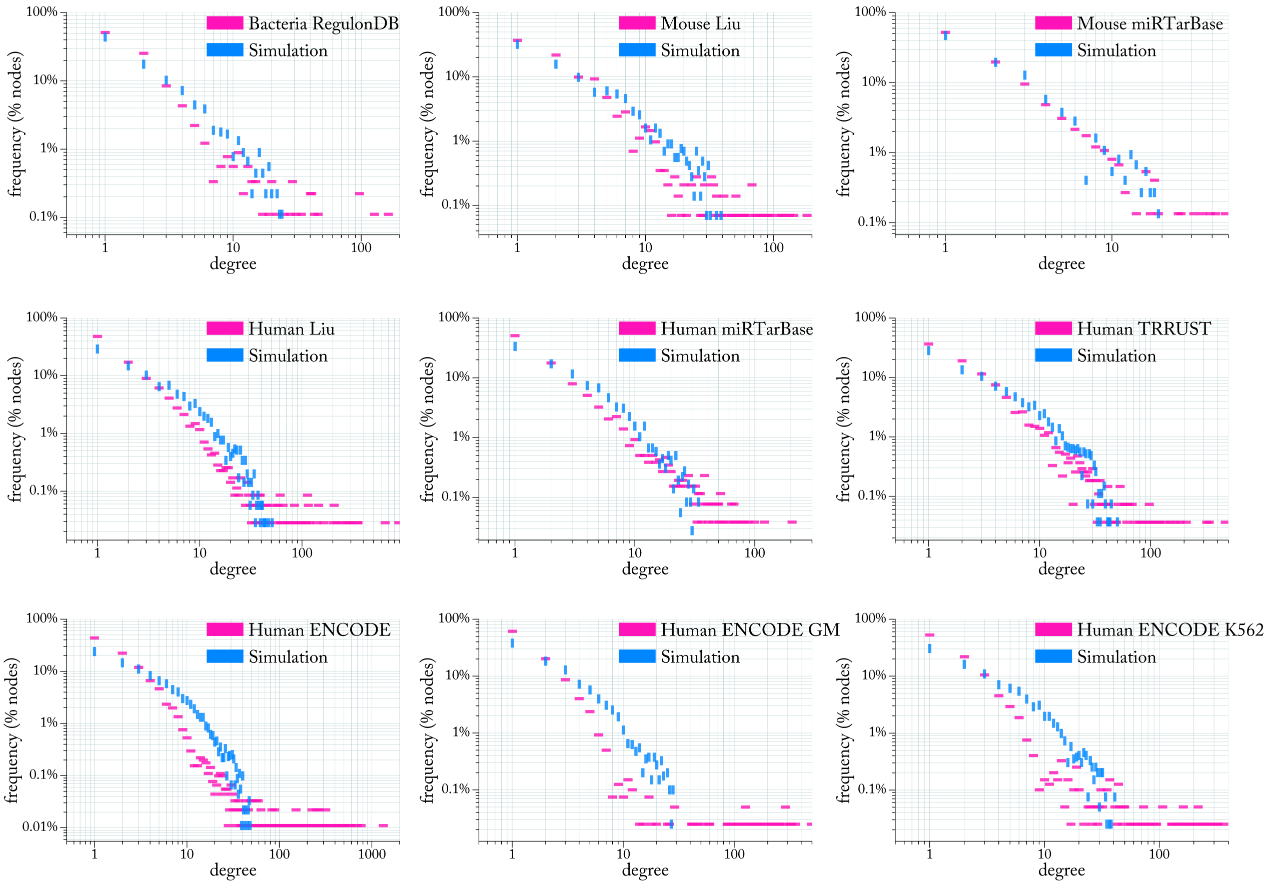


Figure 14: Evolving synthetic networks to the same size (number of nodes and edges) as regulatory networks.

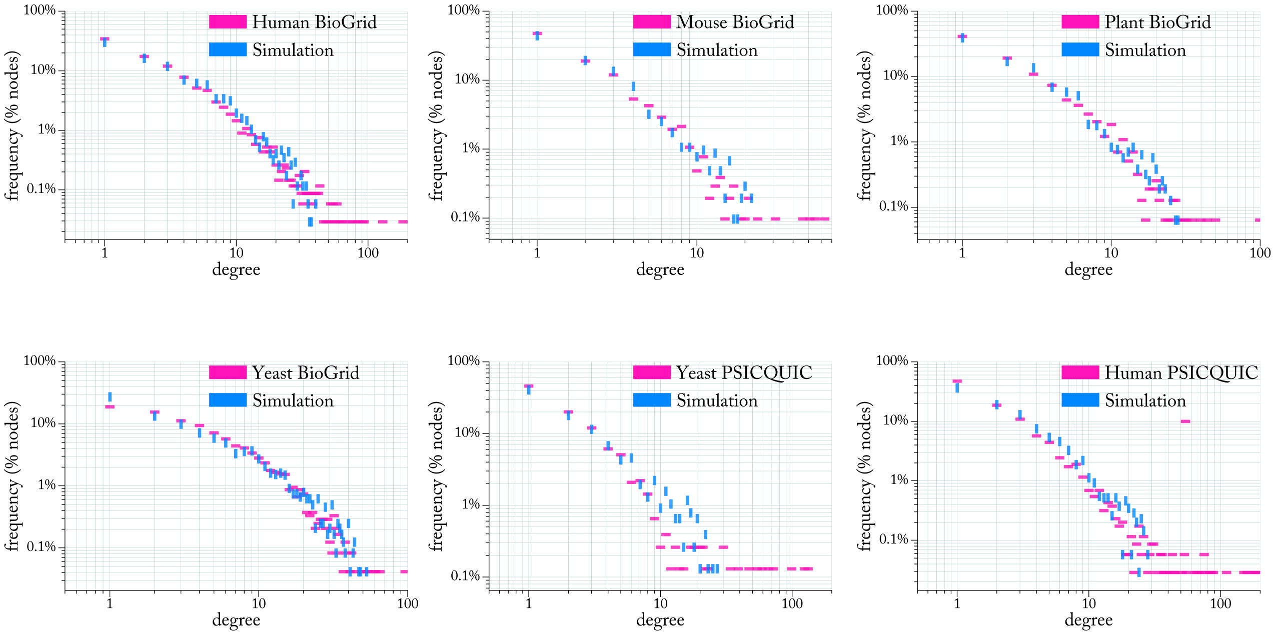


Figure 15: Evolving synthetic networks to the same size (number of nodes and edges) as database-sourced networks.

### 11 Simulated adaptation

In these experiments, the simulated network evolution algorithm (see Figure SI 12) starts with networks that have a number of nodes/edges equal that of a corresponding real MIN. The edges of the seed networks are initially randomly assigned. In each generation, only reassign-edge mutation is carried out (no add-node or add-edge mutations) as opposed to the simulated evolution experiments (Section SI 10). Figures SI 16 and 17 show the degree distribution of the fittest synthetic network (labelled 'Simulation') against that of the corresponding equal-size MIN after 4X and 8X generations of mutate-and-select, respectively, where X equals the number of the nodes in the network. The degree distribution of the initial seed network is labeled 'Seed' in Figures SI 16 and 17. The threshold $t$ of tolerated damaging interactions in the solution is kept at 5% of the sum of all damages in all simulations, as was the case in the simulated evolution experiments of Section 10.

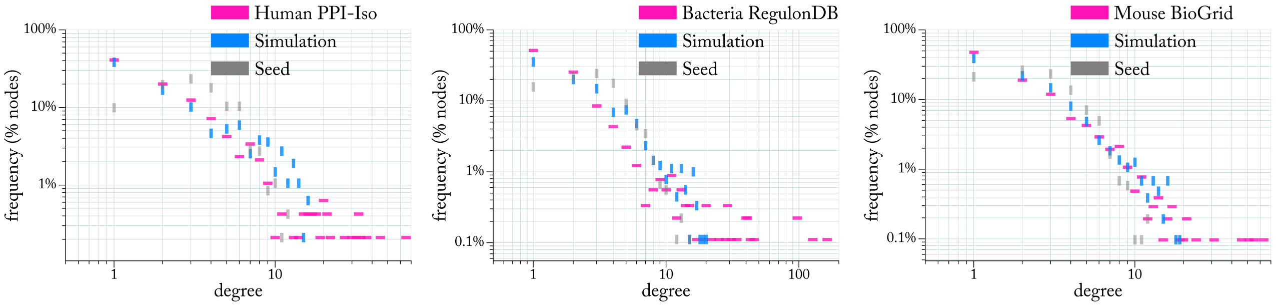


Figure 16: Adapting synthetic networks for 4X generations of mutate-and-select where X is the total number of nodes in the networks. Simulation starts with the a network that has the same number of nodes and edges as the corresponding real MIN, but with edges randomly assigned to nodes. The degree distribution of the synthetic (Simulation) shown here is that of the fittest network after 4X generation of mutate-only simulated adaptation. Increasing the number of generations does not significantly change the degree distribution (see Figure SI 17). The degree distribution of the initial seed network is labeled 'Seed'.

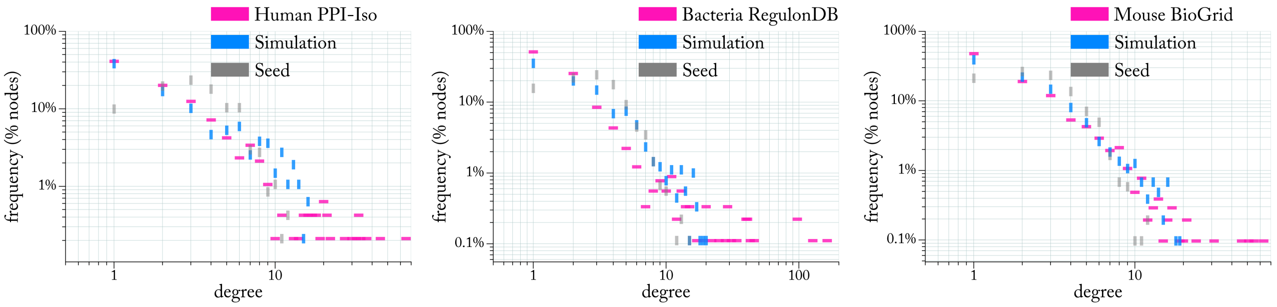


Figure 17: The same simulation as that in Figure SI 16 except that here the simulation is further continued for 4X more generations (hence the total number of generations is 8X the number of nodes in the network). Such increase in number of generations does not significantly change the degree distribution as opposed to simulations terminated after 4X generations (Figure SI 16). The degree distribution of the initial seed network is labeled 'Seed'.
